# Restraining Wnt activation and intestinal tumorigenesis by a Rab35 dependent GTPase relay

**DOI:** 10.1101/2024.02.05.578891

**Authors:** Siamak Redhai, Tianyu Wang, Kim. E Boonekamp, Saskia Reuter, Tümay Capraz, Svenja Leible, Shivohum Bahaguna, Fillip Port, Bojana Pavlović, Michaela Holzem, Roman M. Doll, Niklas Rindtorff, Erica Valentini, Barbara Schmitt, Karsten Richter, Ulrike Engel, Wolfgang Huber, Michael Boutros

## Abstract

Maintenance of homeostatic processes ensure curtailment of intestinal tumorigenesis. Inactivating mutations to Adenomatous Polyposis Coli (*Apc*) result in aberrantly activated Wnt signalling and initiates colorectal cancer (CRC) in approx. 80% of cases, yet our understanding of the subcellular mechanisms that modulate dysregulated pathway activity is limited. Here, using a conditional *in vivo* genetic screen, we identify Rab35 GTPase as a novel tumour suppressor that modulates regional Wnt activity after loss of *Apc* in progenitor cells. Single cell analysis revealed that progenitor cells respond to *Apc* depletion by increasing the expression of a GTPase activating protein, which we named *blackbelt*, and triggering Rab35 disassociation from the plasma membrane. Mechanistically, we demonstrate that Rab35 controls the localisation and activation of the Rho GTPase, Cdc42, which functions as a relay to regulate JNK signalling. This in turn tunes the Wnt pathway upstream of β-catenin to direct proliferation and differentiation of progenitor cells. Importantly, we show that maintaining active JNK signalling is important for the propagation of *Apc* mutant mouse colon organoids. Our findings highlight a novel GTPase cascade that sustains aberrant Wnt activity in specific segments of the intestine and provides impetus to therapeutically exploit this pathway to target CRC.

## MAIN

Precise control over the activity of intestinal stem cells is crucial for maintaining homeostasis and disruptions to these events can lead to numerous gastrointestinal diseases [1]. Colorectal cancer (CRC) is a leading cause of cancer related deaths. Inactivating mutations to the tumour suppressor gene *Adenomatous polyposis coli* (*Apc*) are considered to be an early event that initiates CRC in approx. 80% of cases [2]. APC is a component of the Wnt signalling pathway and has several roles including inhibiting β-catenin, which functions as a transcriptional coactivator in the nucleus and is a component of the adherens junction [3, 4]. The loss of *Apc* function constitutively activates Wnt signalling to drive proliferation and provide cells with a competitive advantage over their surrounding wild type cells, which initiates tumour formation [5–8]. Therefore, understanding how cellular processes are rewired when *Apc* is perturbed may uncover novel pathways that could serve as therapeutic targets.

Endomembranes are increasingly recognised as sites where signals are initiated, terminated or modulated [9–11]. Conversely, signalling activity can also change the flow of material through subcellular compartments by altering membrane trafficking processes [12]. Therefore, this bi-directional cross talk is essential for finely tuning pathway activity to meet cellular demands. Important regulators of membrane trafficking are the Rab and Rho family of GTPases, which are involved in cell polarity, protein secretion, receptor recycling, and cytoskeletal regulation, thereby modulating cell signalling [13–17]. They carry out diverse functions through recruiting a variety of effector proteins in their active GTP bound state [18, 19]. The GTP-GDP cycle of Rab and Rho GTPases are regulated by guanine nucleotide exchange factors (GEFs) and GTPase activating proteins (GAPs) [20, 21]. However, the function of Rab and Rho GTPases and their crosstalk in the intestine are not well understood and many of the subcellular processes required to maintain homeostatic balance in intestinal stem cells remain poorly characterised, despite their recently reported importance in regulating signalling pathway activity [22].

The intestine of *Drosophila melanogaster* has emerged as a powerful *in vivo* model for dissecting the basic molecular mechanisms that are involved in maintaining homeostasis, many of which have been shown to be conserved in mammalian systems [23, 24]. The intestinal epithelium of flies is regionalised at multiple levels and is composed of differentiated enterocytes (ECs), enteroendocrine cells (EEs) and *esg^+^* progenitor cells, which can be divided into intestinal stem cells (ISCs), enteroblasts (EBs) and enteroendocrine progenitors (EEPs) [25–28]. Progenitor cells are able to replenish different cell types in the epithelium and provide an excellent model to investigate the basic cellular and subcellular processes that are involved in maintaining homeostasis [29].

Here, we show that Wnt activation in the intestine is regionalised, occurring predominantly in the anterior segment of the midgut. Using an *in vivo* modifier screen in the *Drosophila* intestine, we identified that *Rab35* modulates regional Wnt signalling when *Apc* is lost in progenitor cells. Using single cell analysis upon *Apc* depletion in the intestine, we show that the expression of a putative GAP, which we named *blackbelt*, is increased in progenitor cells (ISB/EBs), while functional experiments demonstrate its importance for Wnt activation. Mechanistically, we show that loss of *Apc* reduces Rab35 on the plasma membrane of progenitor cells, resulting in Cdc42 activation and membrane translocation. This Rab35-Cdc42 GTPase relay regulates JNK signalling, which acts as a downstream pathway to tune Wnt activity in order to maintain epithelial homeostasis, a pathway we show to be conserved in mouse colon organoids.

## RESULTS

### *Rab35* is a tumour suppressor that acts downstream of *Apc* to regulate progenitor proliferation and lifespan of flies

To investigate how membrane trafficking regulators influence progenitor proliferation when *Apc* is depleted, we selected genes encoding proteins with membrane trafficking functions to screen based on their misexpression in colorectal cancer patients [2] or that were previously associated with regulating signalling cascades. We generated a fly line that inducibly marks progenitor cells with *GFP* and enables *RNAi* to silence *Apc* (*Apc^RNAi^*) in this cell type using the *esg^TS^* driver (*esg-Gal4, tub-Gal80^TS^>GFP*) (Fig. 1a). Previously, it has been shown that loss of *Apc* in the fly intestine induces progenitor proliferation [30–32]. We confirmed this by using *Apc^RNAi^* or with newly generated *gRNAs* targeting *Apc* (*Apc^gRNAx2^*) (Extended data Fig. 1a-c). We introduced a scoring system to assess how genetic interactions between *Apc^RNAi^* and *gene-X* differed from flies expressing *Apc^RNAi^* alone. Genetic interactions that enhanced progenitor proliferation were given a positive score and those that suppressed progenitor numbers were given a negative score. In total, we screened 121 lines covering 63 genes, giving an average of two lines per gene, often including both knockdown and overexpression of targeted candidates (Fig. 1b, see Supplementary Table 1 for Fly ID).

**Fig. 1.**
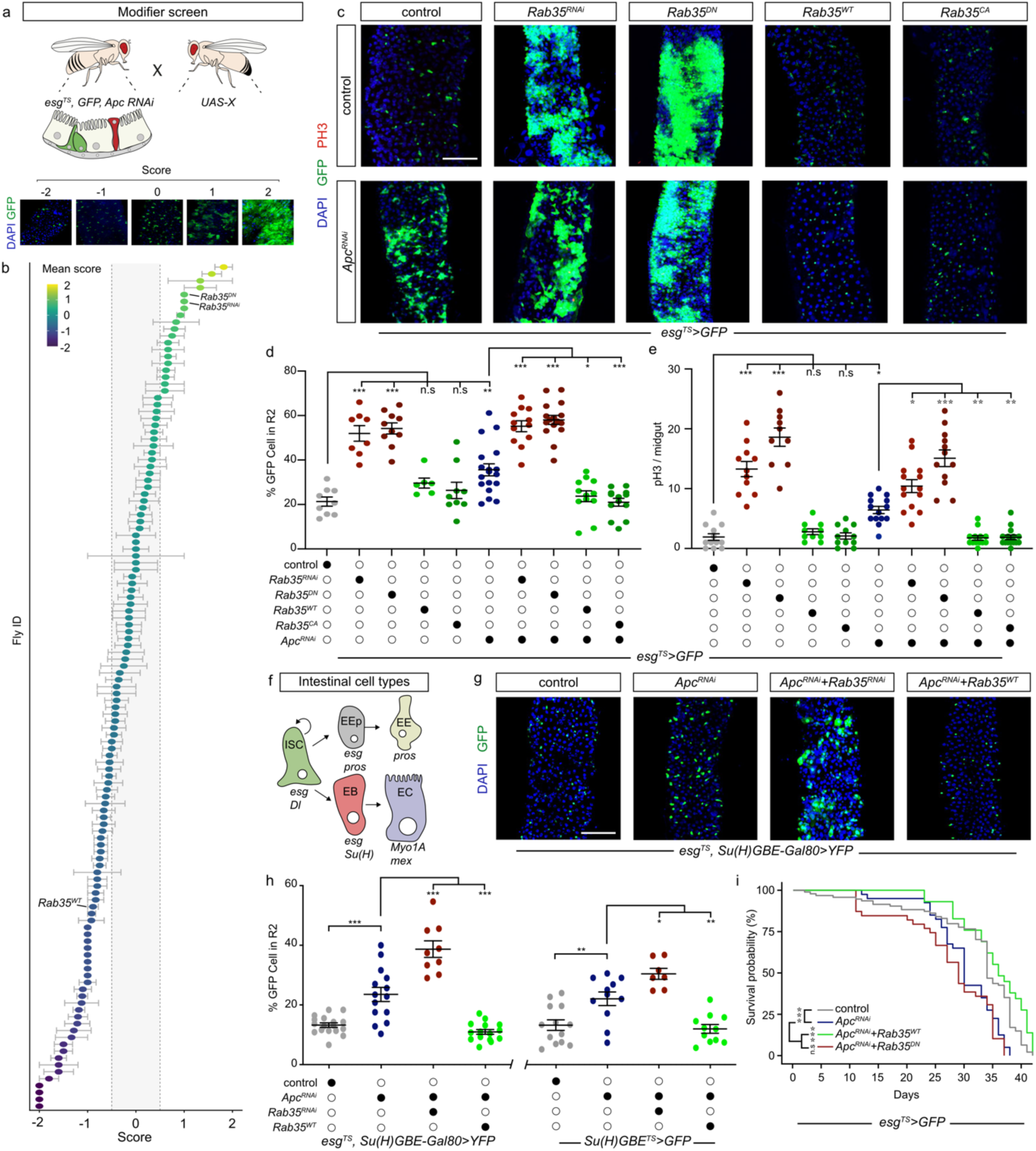
*Rab35* is a tumour suppressor that acts downstream of *Apc* to regulate progenitor proliferation and lifespan. a) Schematic of the *in vivo* modifier screen in this study along with the scoring system used to identify novel modulators. b) We targeted membrane trafficking regulators and scored whether they enhanced (positive score) or supressed (negative score) intestinal progenitor proliferation when *Apc* was knocked down using the *esg^TS^* driver, see supplementary table 1 for Fly ID names. *Rab35* was identified as having both of these properties. Grey zone highlights candidates that were classified as not having a modifying effect. c) *Rab35* functions as a tumour suppressor under homeostasis and can modulate progenitor proliferation when *Apc* is lost in progenitor cells. Mitotically active cells are highlighted with pH3 in red. d) Quantification of *GFP^+^* cells in the R2 region. e) Quantification of pH3 positive cells throughout the entire intestinal epithelium. f) Schematic of intestinal cell types in *Drosophila melanogaster* along with canonical markers. g) Representative images of cell type specific targeting of *Apc* and *Rab35* in ISCs. h) Quantification of *GFP^+^* cells when *Apc* and *Rab35* are targeted in either ISCs only (*esg^TS^, Su(H)GBE-Gal80>GFP*) or EBs only (*Su(H)GBE^TS^>GFP*). i) Overexpressing *Rab35* can rescue the decline in lifespan of flies expressing *Apc RNAi* in progenitor cells. All Graphs represent the mean with standard error of the mean. One-way ANOVA test with Tukey post hoc comparison was used for all graphs except for (i) where a logrank test was used. \**P* < 0.05, \*\**P* < 0.01, \*\*\**P* < 0.001. Scale bar for all images 100 μm.

In *Apc^RNAi^* expressing progenitor cells, *Rab35* was identified as having a suppressive effect on proliferation when its *YFP*-tagged wild type (*Rab35^WT^*) or constitutively active (*Rab35^CA^*) form was expressed. Conversely, *Rab35* enhanced proliferation when its function is perturbed by *RNAi* silencing (*Rab35^RNAi^*) or with a dominant negative *YFP*-tagged mutant (*Rab35^DN^*) (Fig. 1c, d). Consistently, we saw the same trend when we quantified the number of phospho-histone H3 (pH3) positive mitotically active cells throughout the intestine (Fig. 1e). To characterise the role of *Rab35* during homeostasis, we expressed *Rab35^RNAi^* or *Rab35^DN^*alone in progenitor cells and observed that both progenitor proliferation and the fraction of mitotic cells were increased, suggesting that *Rab35* function as a tumour suppressor in the intestine (Fig.1 c-e). We verified this phenotype with Cas9-mediated mutagenesis and new *gRNAs* targeting *Rab35* (*Rab35^gRNAx2^*) (Extended data Fig. 1a-c). Furthermore, overexpression of *Rab35^WT^* or *Rab35^CA^* did not change progenitor cell numbers and the intestine retained its ability to mount an injury response to enteric infection by *Erwinia carotovora carotovora 15* (*Ecc15*) (Extended data Fig. 1d, e).

To test whether the *Apc-Rab35* genetic interaction is observed in other tissues, we performed perturbation experiments in the posterior larval wing disc and looked for morphological changes in the adult wing. Interestingly, expression of *Apc^RNAi^* induced ectopic vein formation at the posterior cross-vein which was phenotypically suppressed by co-expressing *Rab35^CA^* or exacerbated with *Rab35^RNAi^,* suggesting that the *Apc-Rab35* interaction may be a general mechanism for tuning Wnt activity (Extended data Fig. 1f).

Next, we wanted to establish which progenitor proliferation pathways are regulated by *Rab35.* First, we observed that overexpression of the Wnt co-receptor *Arr* (the *Drosophila* homologue of *Lrp6*) increased the number of progenitor cells and co-expressing *Rab35^WT^*reverted this phenotype back to a wild type phenotype (Extended data Fig. 1g, h). Similarly, *Rab35^CA^* was able to block proliferation in cells co-expressing *Apc^RNAi^* and an oncogenic *Ras* mutant (*Ras^V12^*). However, while downregulation of the Notch receptor (*N^RNAi^*) resulted in an expansion of the progenitor population, co-expressing *Rab35^CA^* failed to rescue this phenotype (Extended data Fig. 1g, h). Therefore, *Rab35* regulates specific progenitor proliferation pathways.

To understand the physiological consequences of the *Apc-Rab35* interaction, we first determined which progenitor cell type is responsible for regulating proliferation. Using drivers that target only ISCs (*esg^TS^, Su(H)GBE-Gal80>GFP*) or EBs (*Su(H)GBE^TS^>GFP)* (Fig. 1f), we observed that Rab35 is able to modulate proliferation in *Apc* depleted ISCs and EBs (Fig. 1g, h). Next, we conducted longevity experiments and determined that flies expressing *Apc^RNAi^* using the *esg^TS^* driver had a significant decline in their lifespan, which was fully rescued when *Rab35^WT^* was co-expressed (Fig. 1i). Importantly, during homeostatic conditions, *Rab35^WT^* did not alter the longevity of flies while there was a significant decline with *Rab35^DN^* (Extended data Fig. 2a). To rule out that defects to the intestinal barrier were responsible for the shortening of lifespan, we fed flies food containing blue dye and tested if there was leakage in the abdominal cavity [33]. In all conditions tested, the barrier function of the intestine was preserved (Extended data Fig. 2b). Furthermore, electron micrographs of the intestine showed that junctional integrity was also maintained (Extended data Fig. 2c). In conclusion, our data provide evidence that *Rab35* functions as a tumour suppressor during homeostasis and can modulate progenitor proliferation downstream of *Apc* and *Arr*/*Lrp6*.

**Fig. 2.**
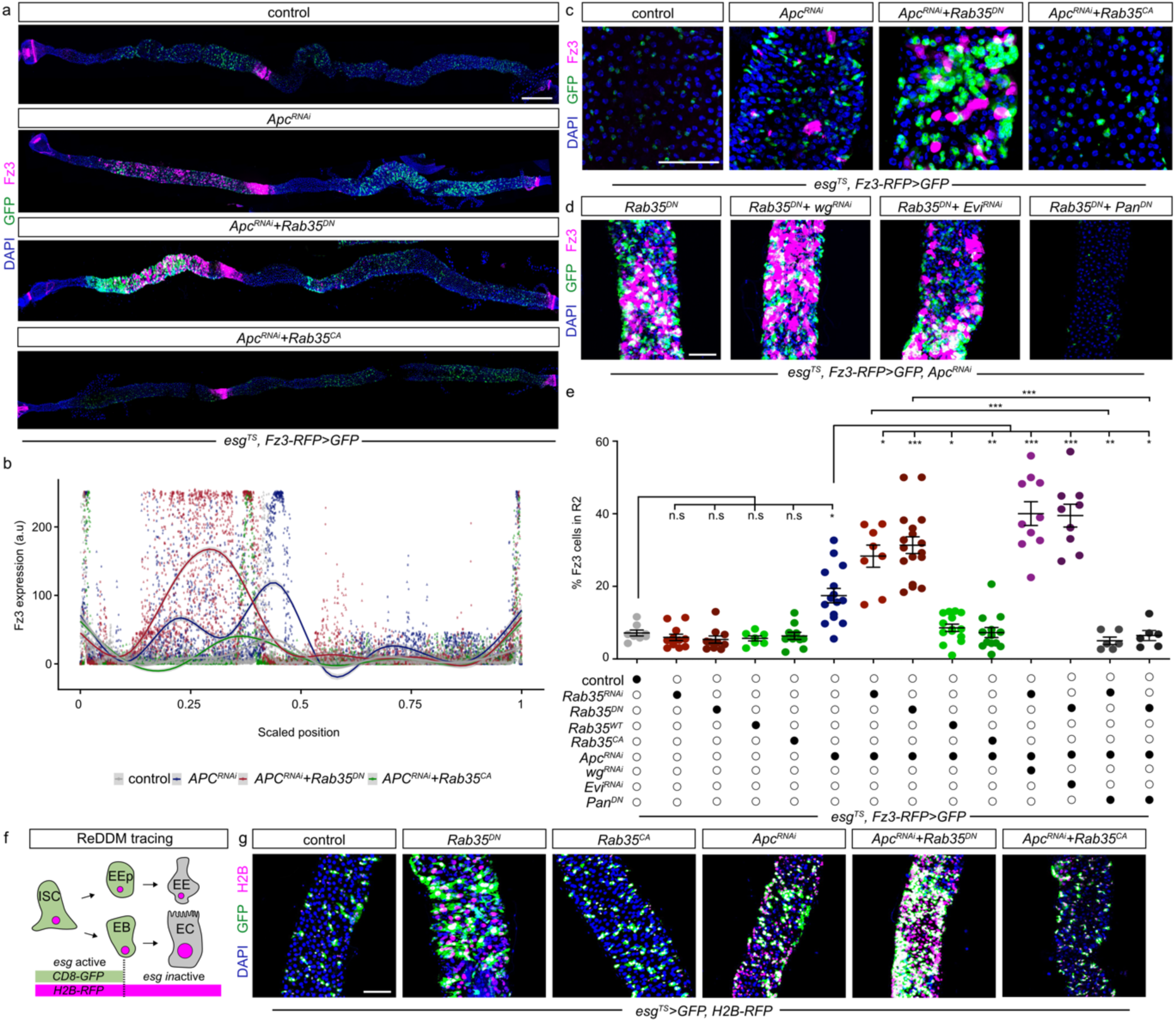
Regional Wnt activation in the intestine is modulated by *Rab35*. a) Wnt activity (*Fz3-RFP*) is prominent at regional boundaries during homeostasis and becomes pronounced in the anterior region when *Apc* is knockdown in all progenitor cells. Expression of a dominant negative *Rab35* mutant (*Rab35^DN^*) exacerbates Wnt activity and progenitor proliferation when *Apc* is knocked down, while a constitutive active *Rab35* mutant (*Rab35^CA^*) rescues these phenotypes. b) Quantification of *Fz3-RFP* along the intestinal epithelium in conditions shown in panel a. Scaled position 0 and 1 corresponds to the proventricular region and MHB boundary, respectively. Lines represent models fitting individual points rather than connect values. c) *Apc* loss in progenitor cells results in cell autonomous and cell non-autonomous Wnt activation (*Fz3-RFP*). The loss of *Rab35* activity (*Rab35^DN^*) can enhance Wnt activation and progenitor proliferation while activation of *Rab35* (*Rab35^CA^*) can rescue this phenotype. d) *Rab35* does not exacerbate Wnt activity and progenitor proliferation through excess wingless (*wg^RNAi^*) or Wnt secretion (*Evi^RNAi^*) but is dependent on *Pan* (*Pan^DN^*). e) Quantification of *Fz3-RFP* in R2 region. f) Schematic of the ReDDM lineage tracing system. g) *Apc* and *Rab35* negatively regulate EC differentiation. *Rab35* activation (*Rab35^CA^*) can reverse EC differentiation when *Apc* is knocked down, while loss of Rab35 accelerates EC differentiation. Graphs represent the mean with standard error of the mean. One-way ANOVA test with Tukey post hoc comparison was used for all graphs. \**P* < 0.05, \*\**P* < 0.01, \*\*\**P* < 0.001. Scale bar for a 200 μm, c, d and g 100 μm.

### *Rab35* regulates regional Wnt activation and progenitor differentiation independently from Wnt secretion

In flies, activation of Wnt signalling results in the expression of Wnt target genes like *Fz3*, *Nkd*, and *Notum* [34]. To gain insight into how *Rab35* regulates Wnt activity when *Apc* is depleted, we monitored *Fz3* using a fluorescently tagged reporter line (*Fz3-RFP*) [35]. Under homeostatic conditions, Wnt activity is highest at compartment boundaries (Fig. 2a, b) [30]. Remarkably, when *Apc* is silenced using the *esg^TS^* driver, Wnt signalling was predominantly activated in the anterior R2 portion of the intestine (Fig. 2a-c). Interestingly, Wnt activation was observed to be both cell-autonomous, in progenitor cells and non-cell-autonomous in the surrounding epithelial cells (Fig. 2c). This was not due to a deleterious effect of *RNAi*, since CRISPR mutagenesis of *Apc1+2^gRNA^*also resulted in a similar anterior Wnt activation pattern and progenitor proliferation (Extended data Fig. 3a-c). To further profile Wnt activity, we generated flip out clones using the *esg^F/O^* driver, which utilises temperature-inducible FLPase expression to activate a constitutive Act>STOP>Gal4 driver by removing the STOP cassette located in between FRT sites [36]. This system drives expression of *GFP* and *UAS-*transgenes in progenitors and their descendants (ECs and EEs). Surprisingly, we saw that in flip out clones of *Apc^RNAi^*, Wnt signalling was not uniformly activated in *GFP^+^*cells, suggesting that the response to *Apc^RNAi^* may be stochastic or depend on cellular state (Extended data Fig. 3d).

**Fig. 3.**
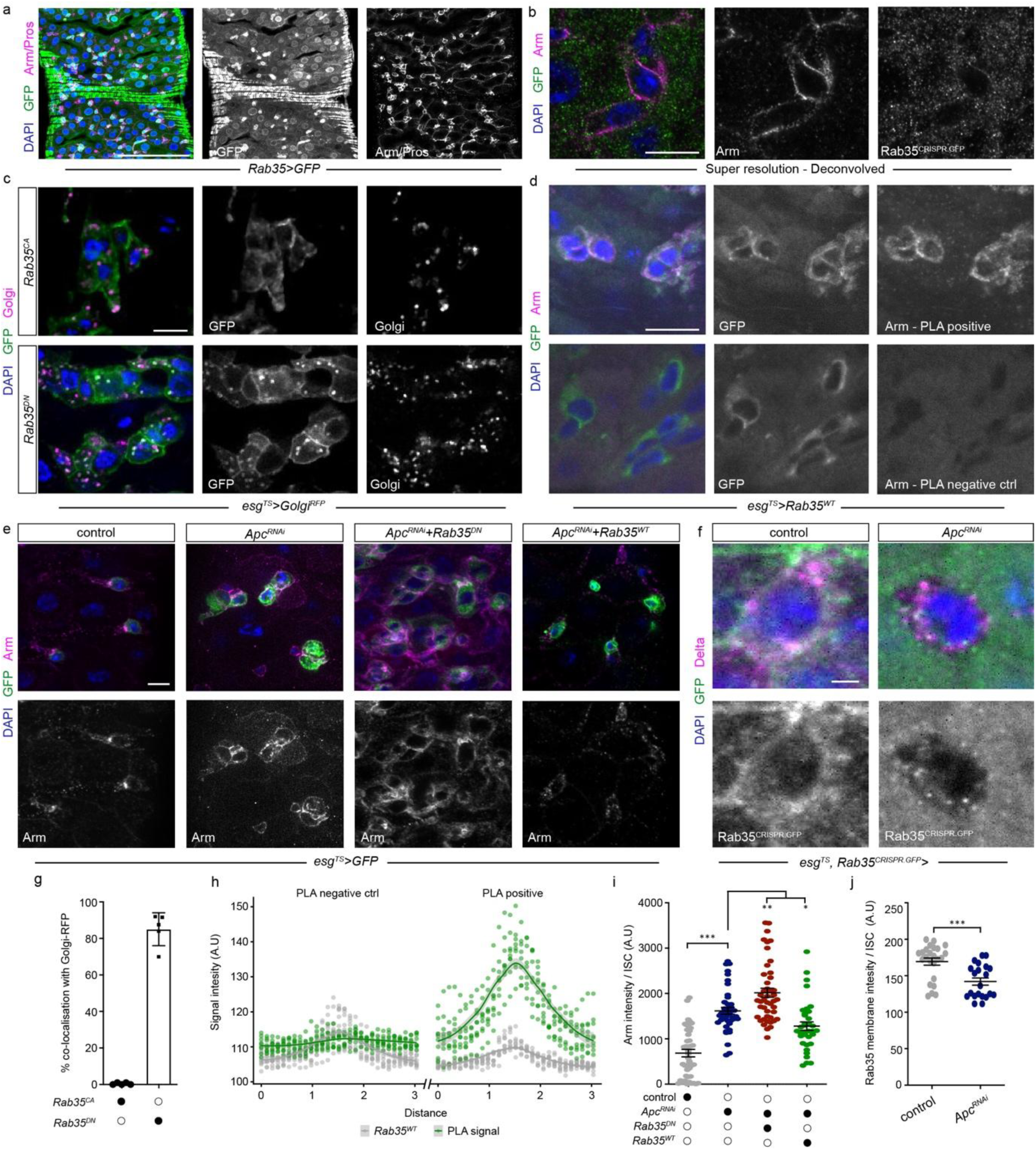
Rab35 co-localises with Arm and regulates its membrane levels. a) *Rab35* expression (*Rab35-Gal4*) is present in all intestinal epithelial cells and is prominent in progenitor cells and the surrounding visceral muscle. b) Super-resolution microscopy reveals endogenous Rab35 is prominent on progenitor cell membrane and co-localises with Arm. c) Rab35 activity directs its intracellular localisation as overexpressing a *YFP*-tagged dominant negative mutant of Rab35 (*Rab35^DN^*) results in its translocation to intracellular puncta found at the Golgi (*Golgi^RFP^*), while no such effect is seen with a *YFP*-tagged wild type version of Rab35 (*Rab35^WT^*). d) Proximity ligation assay demonstrates that Rab35 and Arm are <40nm apart. e) Loss of *Apc* increases membrane Arm in R2, while Rab35 negatively modulates this phenotype. f) Endogenous Rab35 is less prominent on the membrane of Delta^+^ ISCs when *Apc* is knocked down in progenitor cells. g) Quantification of the co-localisation between different *Rab35* constructs with *Golgi^RFP^* in progenitor cells. h) Quantification of PLA signal on the progenitor plasma membrane. i) Quantification of membrane Arm intensity on progenitor cells in R2. j) Quantification of Rab35 membrane intensity on ISCs. Graphs represent the mean with standard error of the mean. One-way ANOVA test with Tukey post hoc comparison was used for all graphs. \**P* < 0.05, \*\**P* < 0.01, \*\*\**P* < 0.001. Scale bar a 100 μm, b, c, e and F 10 μm.

To understand how Rab35 modulates the Wnt pathway, we performed epistasis experiments with *Apc^RNAi^*, which demonstrated that Rab35 is able to regionally enhance or suppress Wnt activity by co-expressing *Apc^RNAi^* with either *Rab35^DN^* or *Rab35^CA^*, respectively. (Fig. 2a-c). Importantly, *Rab35^DN^* or *Rab35^CA^* had no impact on Wnt signalling on its own, suggesting that Rab35 modulates the Wnt pathway only when *Apc* is depleted (Fig. 2e). Rab35 has previously been implicated in the fast recycling pathway [37]. To define the specificity of Rab35 for Wnt regulation, we tested the role of other Rab GTPases previously implicated in the recycling pathway [38]. We found that depletion of Rab4 or Rab11 in progenitor cells expressing *Apc^RNAi^* did not phenocopy the loss of Rab35 (Extended data Fig. 3e-g). Importantly, while knockdown of the early endosomal Rab5 exacerbated progenitor proliferation, it had no impact on Wnt pathway activity (Extended data Fig. 3e-g). These data demonstrate that the *Rab35*-dependent trafficking pathway can function unlike other recycling Rab GTPases in modulating Wnt activity.

Activating mutations to β-catenin have been implicated in colorectal cancer [39]. To determine where *Rab35* functions along the Wnt pathway, we performed genetic interaction experiments with a constitutively active *Arm* (*Arm^CA^*) mutant, the fly homolog of β-catenin. Expression of *Arm^CA^* using the *esg^TS^* driver resulted in a moderate expansion of progenitor cells and a largely cell-autonomous activation of the Wnt pathway (Extended data Fig. 3h-j). Interestingly, co-expression of *Rab35^WT^* or *Rab35^CA^* with *Arm^CA^* had no effect on proliferation or Wnt activity (Extended data Fig. 3h-j). In contrast, co-expression *Arm^CA^* with a dominant negative *Pangolin* (*Pan^DN^*) mutant, which is the main Wnt transcription factor and homolog of the lymphoid-enhancing factor (LEF)/T cell factor (TCF), was able to fully suppress progenitor proliferation and Wnt activity (Extended data Fig. 3h-j). Thus, Rab35 functions downstream of *Apc* but upstream of *Arm* to modulate Wnt signalling and progenitor proliferation.

Since we observed high Wnt signalling in ECs when *Apc^RNAi^*was expressed in progenitor cells, we wanted to test the contribution of Wnt activation in ECs for intestinal homeostasis. Interestingly, silencing *Apc* in ECs using the *Myo^TS^* driver (*Myo1A-Gal4, tub-Gal80^TS^>GFP*) results in anterior Wnt activation (Extended data Fig. 4a, b). However, this was not accompanied by an increase in progenitor proliferation or a decline in the longevity of flies, providing evidence that Wnt activation in ECs is dispensable for these phenotypes (Extended data Fig. 4c, d).

**Fig. 4.**
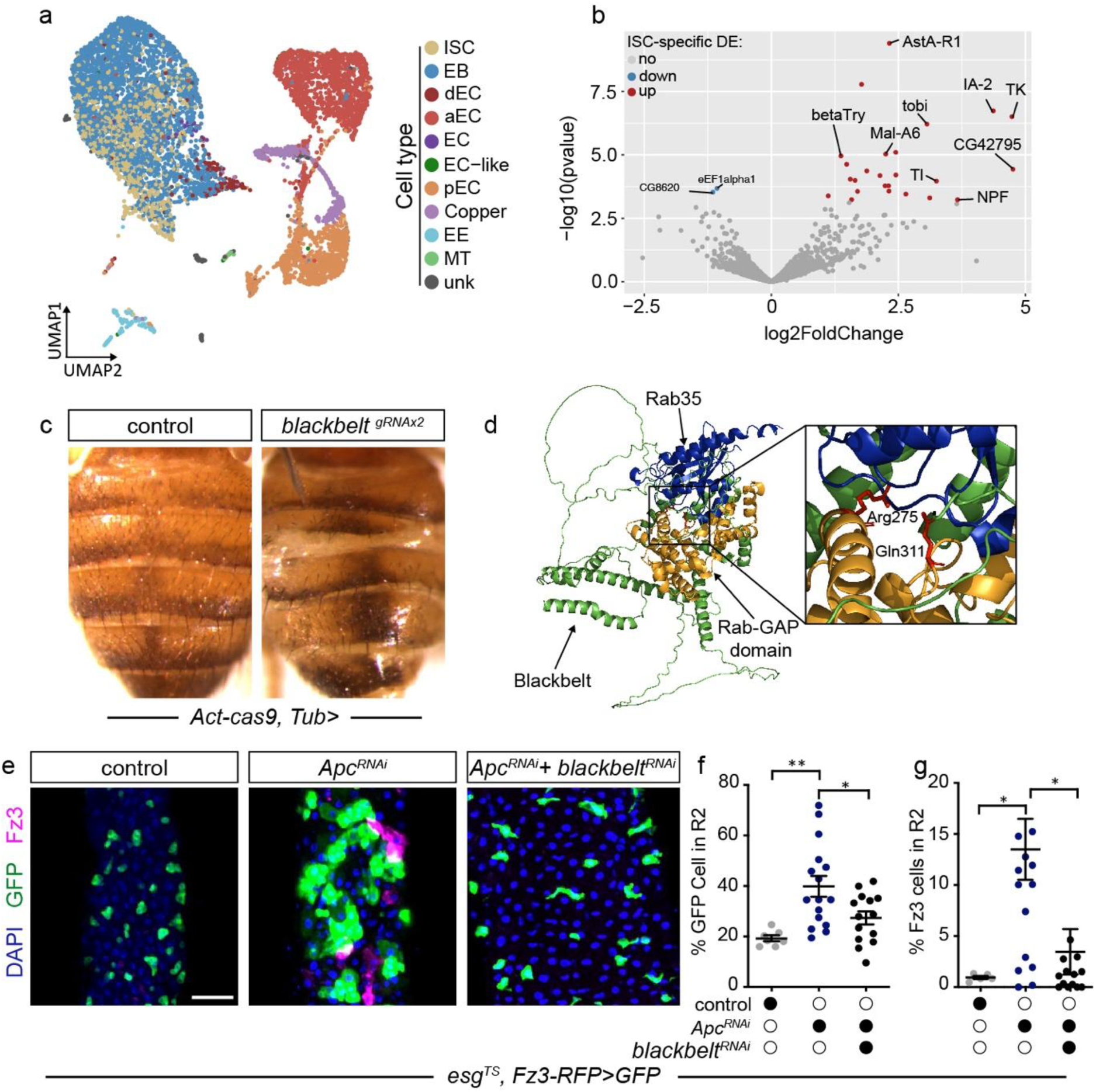
*blackbelt* is a putative GAP and is involved in modulating Wnt signalling. a) scRNA-seq of the intestine during homeostasis and progenitor-specific knockdown of *Apc* was performed – control uMAP and cell types are shown. b) *CG42795* expression is significantly increased in the ISC population when *Apc* is knocked down in progenitor cells using the *esg^TS^* driver. c) *CG42795* was named *blackbelt* due to its loss of function phenotype – CRISPR generated mutant flies (*Act-cas9, Tub> blackbelt^gRNAx2^*) display a prominent black stripe on the abdomen with some missing abdominal cuticles. d) Structural modelling revealed that Blackbelt is a conventional GAP that are defined by their arginine and glutamine residues which interact with Rab35. e) *blackbelt* is required for Wnt activation and progenitor proliferation when *Apc* is knocked down using the *esg^TS^* driver. f) Quantification of *GFP^+^*cells in R2. g) Quantification of *Fz3-RFP* in R2 region. Graphs represent the mean with standard error of the mean. One-way ANOVA test with Tukey post hoc comparison was used for all graphs. \**P* < 0.05, \*\**P* < 0.01, \*\*\**P* < 0.001. Scale bar e 100 μm.

*Rab35* has been implicated in a number of trafficking and secretory processes [40, 41], thus we wanted to investigate if *Rab35* exacerbates Wnt signalling through excess Wnt secretion. To test this, we knocked down the fly Wnt1 homologue, *wingless* (*wg*), or the Wnt secretory cargo receptor *Evi/Wls* in progenitor cells expressing *Rab35^DN^ or Rab35^RNAi^* and *Apc^RNAi^*. As shown in Fig. 2d and e, depletion of *Wg* or *Evi* in this genetic background had no impact on proliferation or Wnt activity [42]. Crucially, we show that expression of *Pan^DN^*can completely suppress these two phenotypes in progenitor cells expressing *Rab35^DN^ or Rab35^RNAi^* and *Apc^RNAi^*, suggesting that Rab35 does not enhance Wnt activity through excessive Wnt secretion but does depend on *Pan* to exert its function (Fig. 2d, e).

Wnt signalling has previously been shown to regulate intestinal stem cell fate decisions [43]. To study this, we used the ReDDM lineage tracing system which marks progenitor cells in both *mCD8-GFP* and *H2B-RFP* and their differentiated descendants with *H2B-RFP* only (Fig. 2f) [44]. Using the *esg^ReDDM^*, we found that both *Rab35* and *Apc* act as negative regulators of EC fate, since their loss increased the turnover of this cell type in the intestine (Fig. 2g). Consistently, while *Rab35^CA^* expression had limited impact on epithelial turnover during homeostasis, it was able to suppress EC fate in flies expressing *Apc^RNAi^*, while *Rab35^DN^* accelerated the EC differentiation program (Fig. 2G). In conclusion, *Rab35* is able to steer the differentiation trajectory of Wnt activated progenitor cells.

### *Rab35* localisation and regulation of membrane-bound *Arm*

To understand how *Rab35* modulates Wnt signalling, we studied its expression and localisation in the intestine. Using a transcriptional reporter line that mimics endogenous *Rab35* expression, we observed that *Rab35* was prominent in the progenitor population, but also in EEs, ECs and the surrounding visceral muscle (Fig. 3a). Super-resolution microscopy revealed that endogenously tagged Rab35^CRISPR.GFP^ co-localised with Arm on the plasma membrane of progenitor cells (Fig. 3b). Similarly, overexpression of *YFP*-tagged *Rab35^WT^* also co-localised with Arm on progenitor cell membranes (Extended data Fig. 5a).

**Fig. 5.**
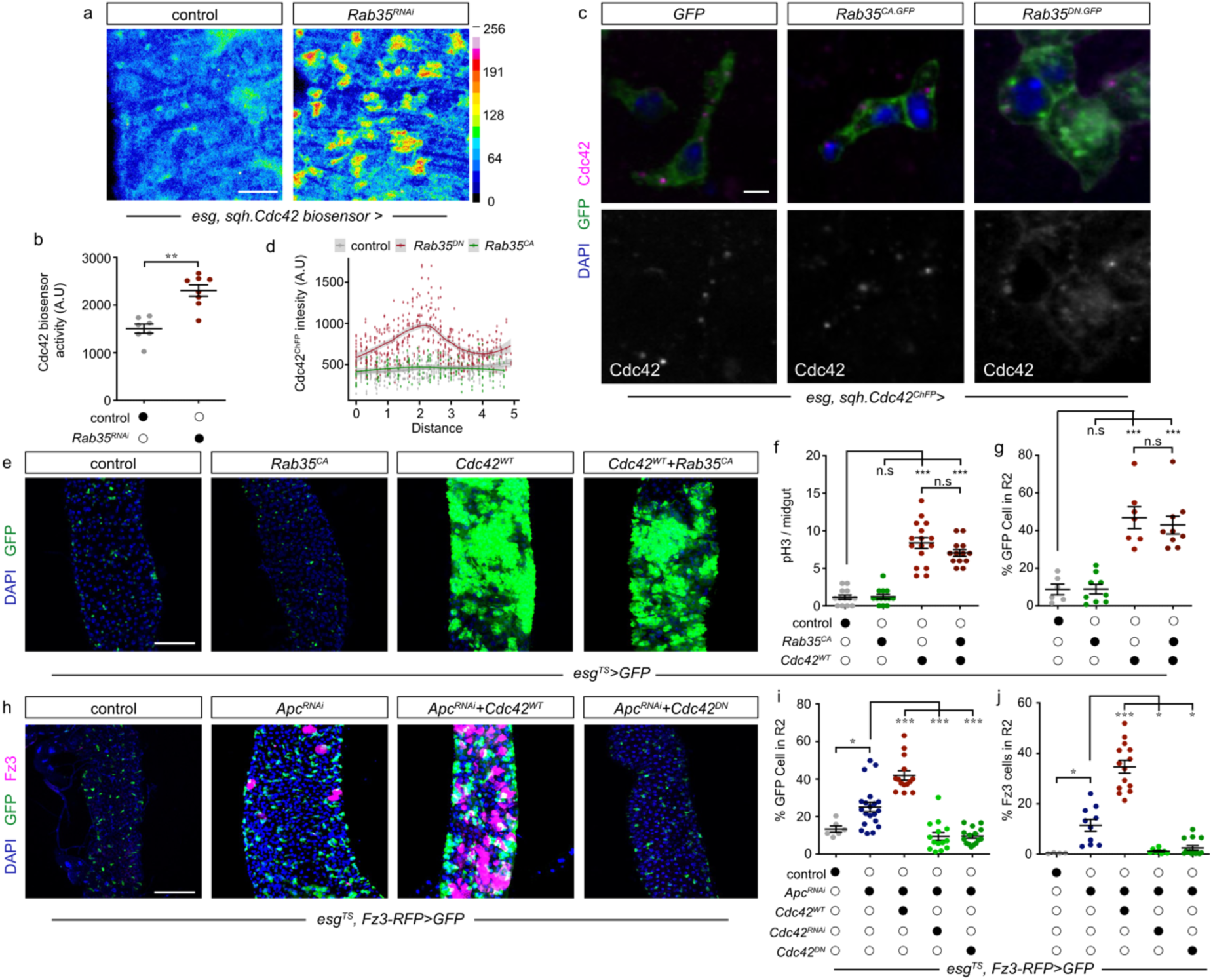
*Rab35* activates *Cdc42* which positively regulates Wnt signalling and progenitor proliferation. a) *Rab35* knockdown in progenitor cells using the *esg* driver increases Cdc42 activity as measured by a *Cdc42* biosensor. b) Quantification of Cdc42 biosensor in progenitor cells. c) Cdc42 is normally present as puncta in progenitor cells during homeostasis and its membrane localisation is negatively regulated by *Rab35*. d) Quantification of Cdc42 on progenitor cell membrane. e) Overexpression of wild type *Cdc42* increases progenitor population and this phenotype cannot be rescued by expressing *Rab35^CA^*. f) Quantification of midgut mitosis (pH3) throughout the whole intestine. g) Quantification of *GFP^+^* cells in the R2 region. h) Overexpressing wild type *Cdc42* enhances progenitor proliferation and Wnt activation when *Apc* is knocked down using the *esg^TS^* driver, while inactivating *Cdc42* (*Cdc42^DN^*) can recues these phenotypes. i) Quantification of *GFP^+^* cells in the R2 region. j) Quantification of Wnt activity (*Fz3-RFP*) in the R2 region. Graphs represent the mean with standard error of the mean. One-way ANOVA test with Tukey post hoc comparison was used for all graphs except for b where a Mann-Whitney test was used. \**P* < 0.05, \*\**P* < 0.01, \*\*\**P* < 0.001. Scale bar a 50 μm, c 2 μm, e and h 100 μm.

*Rab35* exists in an active GTP and inactive GDP bound state [45]. To reveal how Rab35 activity influences its localisation, we utilised *YFP*-tagged constitutively active *Rab35^CA^*and dominant negative *Rab35^DN^* mutants. Interestingly, we found that *Rab35^DN^*, while present on the plasma membrane, also appeared on intracellular puncta located at the Golgi, whereas active *Rab35^CA^* had limited overlap with the Golgi and was predominately plasma membrane bound (Fig. 3c, g). Therefore, the subcellular localisation of Rab35 is determined by its activity state. To investigate if Rab35 and Arm could potentially interact on the plasma membrane, we used *in situ* proximity ligation assay (PLA), which is capable of detecting protein-protein interactions that are within a 40 nm distance [46]. Indeed, we detected positive PLA signal corresponding to proximal *Rab35^WT^*and Arm on the progenitor cell membrane, which was absent in negative controls (Fig. 3d, h).

As well as translocating to the nucleus upon Wnt activation, β-catenin has been shown to increase on the cell membrane when Wnt signalling is activated [47–49]. In flies, detecting nuclear Arm is difficult unless nuclear export inhibitors are used [50], however, this did not work in the intestine. To understand if *Apc-Rab35* regulates Arm levels, we studied Arm membrane intensity on progenitor cells. Using an endogenously GFP-tagged Arm, we show that expression of *Apc^RNAi^* increased Arm on the progenitor cell membrane (Extended Data Fig. 5b, c). Similarly, we observed the same phenotype when using an antibody raised against Arm (Fig. 3e, i). Notably, co-expression of *Rab35^WT^*with *Apc^RNAi^* resulted in a reduction of membrane Arm when compared to *Apc^RNAi^*, while *Rab35^DN^* significantly increased membrane Arm (Fig. 3e, i). This suggests that membrane-bound Arm, in part, corresponds to Wnt activity and that *Rab35* can regulate its abundance.

To understand how *Apc* may regulate Rab35, we studied the localisation of endogenous Rab35^CRISPR.GFP^ during homeostasis and *Apc* knockdown. Surprisingly, we observed that while Rab35^CRISPR.GFP^ is predominantly membrane-bound on ISCs, its membrane association decreased significantly under *Apc^RNAi^* expression (Fig. 3f, j). Since Rab35 activity is known to regulate its localisation, we conclude that *Apc^RNAi^* may decrease Rab35 activity and lead to its dissociation from the plasma membrane.

To determine the trafficking events that are regulated by *Rab35*, we focused on the localisation of GFP-tagged human transferrin receptor (hTfR^GFP^) which is an established recycling cargo [37]. Interestingly, loss of function of either *Apc* or *Rab35* resulted in heterologous expression of hTfR^GFP^ since there were significantly more intracellular hTfR^GFP^ puncta (Extended Data Fig. 5d, e). This indicates that both *Apc* and *Rab35* are involved in some aspects of hTfR^GFP^ recycling. Using electron microscopy, we show that neither expression of *Apc^RNAi^* nor co-expression of *Apc^RNAi^* with *Rab35^DN^* significantly altered the Golgi architecture in progenitor cells (Extended Data Fig. 5f), suggesting that the increase in membrane Arm in this condition is not due to defects to this organelle but may rather result from misregulation of recycling events.

### *blackbelt* is a putative GAP that modulates Wnt signalling

To gain insights into the transcriptional events upon loss of *Apc* in different cell types, we next performed single-cell RNA-sequencing (scRNA-seq) during homeostasis and upon progenitor specific expression of *Apc^RNAi^*. scRNA-seq experiments were run in duplicates and we recovered 14,737 cells in total after quality control (control: 9136, *Apc^RNAi^*: 5601) (Fig. 4a and Extended Data Fig. 6a). Clustering analysis revealed 9 major cell types consistent with previous studies [51, 52]. Differential gene expression analysis highlighted *CG42795* as being significantly upregulated only in the progenitor population (ISC, EB, ISC/EB) when *Apc* was silenced in these cell types (Fig. 4a, b and Extended Data Fig. 6b-e). *CG42795* is an uncharacterised gene, but is predicted to enable GTPase activity, which is characteristic of Rab GAP proteins that negatively regulate the activity of Rab GTPases [53]. Evolutionary analysis of *CG42795* across different species highlighted a conserved TBC (Tre2/Bub2/Cdc16) domain and a large extension at the C-terminus that is present only in *Drosophila* (Extended Data Fig. 7a, b). Next, we generated *CG42795* knockout flies using CRISPR mutagenesis (Extended Data Fig. 7c). Mutant flies were viable and presented with a pronounced stripe of abdominal pigmentation and some missing abdominal cuticles (Fig. 4c and Extended Data Fig. 7d). We therefore named this gene *blackbelt*, owing to its loss of function phenotype.

Structural modelling of Blackbelt highlighted conserved and well predicted domains, however, the long *Drosophila* specific C-terminal region was not modelled reliably (Fig. 5d). Structural predictions between Blackbelt and Rab35 revealed a potential interaction of these two proteins in the Rab-TBC-GAP domain (Fig. 5d). The interaction between these two proteins involves two catalytic amino acids, called the R (Arg275) and Q (Gln311) finger [54, 55]. Using ColabFold, both amino acids, Arg275 and Gln311, were detected in the predicted interaction domain of Blackbelt with Rab35, making Blackbelt a conventional GAP protein. We therefore hypothesised that *blackbelt* may be involved in regulating Wnt signalling in the intestine. Indeed, co-expression of *blackbelt^RNAi^* with *Apc^RNAi^* decreases progenitor proliferation and Wnt signalling, mimicking the phenotype observed when combining *Rab35^CA^* and *Apc^RNAi^* (Fig. 4e-g). This data supports that Blackbelt is important for progenitor proliferation and Wnt activation.

### *Cdc42* activity and localisation is negatively regulated by *Rab35*

We next wanted to understand the Rab35-dependent downstream mechanisms that modulate Wnt signalling and progenitor proliferation. The cross talk between Rab and Rho GTPases are poorly understood in the intestine, especially in the context of Wnt signalling. Therefore, we investigated this by focusing on the Rho GTPase, *Cdc42*, since its activity has been reported to be regulated by *Rab35* during neurite outgrowth *in vitro* [56]. We first addressed whether *Rab35* regulates *Cdc42* activity in the intestine by using a GFP-tagged p21-Rho binding domain of WASp to monitor active Cdc42 GTPase – a Cdc42 biosensor [57, 58]. Knockdown of *Rab35* resulted in an enrichment of Cdc42 biosensor activity in progenitor cells (Fig. 5a, b and Extended Data Fig. 8a). *Cdc42* has been reported to be active at the cell periphery [59, 60]. Consistent with this, localisation of Cdc42^ChFP^ changed from punctate-like structures in control or *Rab35^CA^* condition to associating with the plasma membrane when *Rab35^DN^* was expressed in progenitor cells (Fig. 5c, d). Thus, Rab35 can determine the activation and localisation of Cdc42.

Since *Rab35* was able to activate Cdc42, we investigated the consequence of *Cdc42* gain of function in the intestine. Overexpression of a wild type form of *Cdc42*, *Cdc42^WT^*, using the *esg^TS^* driver led to a massive expansion of progenitor and mitotically active cells (Fig.5e-g). Moreover, lineage tracing experiments highlighted that *Cdc42* positively regulates EC differentiation since overexpression of *Cdc42^WT^* with *esg^ReDDM^* led to an increase in EC turnover (Extended Data Fig. 8b). Consistent with *Cdc42* being downstream of *Rab35*, expression of *Rab35^CA^* was not able to suppress *Cdc42^WT^*-induced progenitor proliferation or EC differentiation (Figure. 5e-g and data not shown). Therefore, Cdc42 positively regulates progenitor proliferation and EC turnover.

Our current data supports that *Rab35* regulates *Cdc42,* which in turn controls the proliferative capacity of progenitor cells. If Cdc42 is a downstream target of Rab35, we expect that it could also modulate Wnt signalling. To this end, we performed genetic interaction experiments between *Cdc42* and *Apc*. Interestingly, co-expression of *Cdc42^WT^* with *Apc^RNAi^* led to significantly higher Wnt activity and progenitor proliferation when compared to *Apc^RNAi^* alone (Fig. 5h-j). Conversely, we could revert *Apc^RNAi^*-induced Wnt activation and progenitor proliferation back to wild type levels when *Cdc42* was silenced with *RNAi* or by co-expressing a dominant negative *Cdc42* mutant (*Cdc42^DN^*) (Fig. 5h-j). Taken together, we show that *Cdc42* is necessary and sufficient to modulate Wnt activity in the absence of *Apc*.

### JNK signalling has a conserved role in modulating progenitor proliferation and Wnt signalling in the intestine

We next asked which signalling cascades downstream of the *Apc-Rab35-Cdc42* axis modulate the Wnt pathway and progenitor proliferation. Previous reports indicate that *Cdc42* regulates JNK signalling *in vitro*, but the role of this interaction in the intestine has not been studied before *in vivo* [61]. To investigate whether JNK signalling is involved in the *Apc-Rab35-Cdc42* axis, we examined the expression of a JNK reporter, *puc-lacZ*. Interestingly, expression of either *Rab35^DN^* or *Cdc42^WT^* in progenitors resulted in widespread JNK activation both in progenitor cells and surrounding ECs, while a modest increase in JNK signalling was observed with *Apc^RNAi^* (Fig. 6a-b). Consistent with *Rab35^WT^* having limited effects on homeostatic processes, its overexpression did not alter the activity of JNK signalling (Fig. 6a-b). Next, we assessed what contributions JNK signalling has during progenitor proliferation. Epistasis experiments revealed that JNK signalling acts downstream of *Rab35* and *Cdc42*, since JNK pathway suppression by a dominant negative form of *Bsk* (*Bsk^DN^*) blocked *Rab35^DN^* or *Cdc42^WT^*-induced progenitor proliferation (Fig. 6c-d). We hypothesised that if JNK signalling is downstream of *Rab35* and *Cdc42*, then its activation should also increase EC differentiation. Indeed, using the *esg^ReDDM^* we demonstrate JNK activation, by *RNAi*-mediated silencing of the negative regulator *puc*, accelerated the production of ECs (Extended Data Fig. 9a). In conclusion, our data supports that the JNK pathway is a downstream target of the Rab35-Cdc42 GTPase axis.

**Fig. 6.**
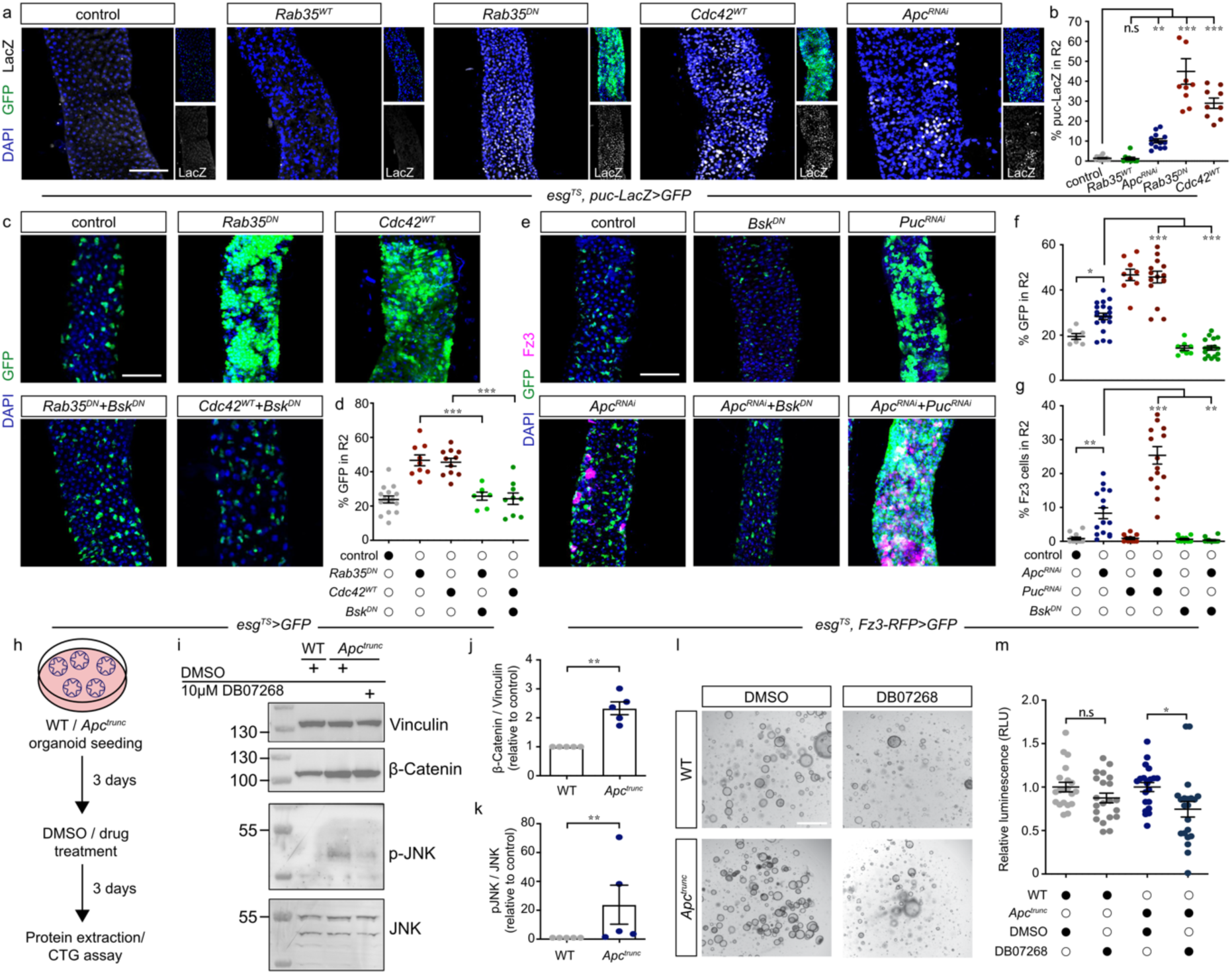
JNK signalling downstream of *Rab35* and *Cdc42* modulates Wnt activity and progenitor proliferation. a) *Rab35* and *Apc* negatively regulate JNK activity, while *Cdc42* positively regulates this pathway as measured by *puc-LacZ* expression in the R2 region. b) Quantification of *puc-lacZ* positive nuclei in R2 region. c) Blocking JNK signalling by expression of a dominant negative mutant form of *Bsk* (*Bsk^DN^*) can rescue progenitor cell proliferation when wild type *Cdc42* is overexpressed or *Rab35* activity is lost (*Rab35^DN^*). d) Quantification of *GFP^+^* cells in R2 region. e) Activation of JNK signalling (*puc^RNAi^*) can enhance Wnt activity and progenitor cell proliferation when *Apc* is lost using the *esg^TS^* driver, while blocking JNK signalling (*Bsk^DN^*) can rescue these phenotypes. Notice that manipulating the JNK pathway during homeostasis has no impact on Wnt activity. f) Quantification of *GFP^+^* cells in the R2 region. g) Quantification of Wnt activity (*Fz3-RFP*) in R2 region. h) Schematic of colon organoid experimental procedure. i) Immunoblot in WT and Apc^trunc^ organoids to measure JNK activity and β-catenin. Notice that p-JNK is elevated in Apc^trunc^ organoids and can be reduced by treatment with DB07268. j) Quantification of β-catenin over Vinculin which was used as a loading control. k) Quantification of p-JNK over total JNK. l) Images of WT and Apc^trunc^ organoids before and after treatment with either DMSO or DB07268. m) Quantification of the relative luminescence from a CTG assay. Notice that Apc^trunc^ organoid growth is reduced when treated with DB07268. Graphs represent the mean with standard error of the mean. One-way ANOVA test with Tukey post hoc comparison was used for all graphs except for j, k and m where a Mann-Whitney test was used. \**P* < 0.05, \*\**P* < 0.01, \*\*\**P* < 0.001. Scale bar a, c, e 100 μm, l 200 μm.

To understand the convergence between JNK and Wnt signalling, we investigated their epistatic relationship in the intestine. First, we found that expression of *puc^RNAi^* led to an expansion of progenitor cells, while it had no effect on Wnt activity during homeostasis (Fig. 6e-g). Second, blocking JNK signalling with *Bsk^DN^* did not alter Wnt activity or progenitor numbers during homeostasis (Fig. 6e-g). Thus, JNK signalling does not regulate the Wnt pathway during homeostatic conditions. Interestingly, when Wnt signalling was activated by *Apc^RNAi^*, co-expression of *Bsk^DN^*or *puc^RNAi^* either suppressed Wnt activity and progenitor proliferation or enhanced these two phenotypes, respectively (Fig. 6e-g). This suggests a requirement for JNK signalling during Wnt transduction specifically in the absence of *Apc*. In order to understand where JNK signalling is involved along the Wnt pathway, we tested its synergy with *Arm^CA^*. Interestingly, activation of Wnt signalling by *Arm^CA^* could not be blocked by expressing either *Bsk^DN^*or silencing the JNK positive regulator hemipterous (*hep^RNAi^*) (Extended Data Fig. 9b). Therefore, we conclude that JNK signalling acts as an integration point to transduce the *Rab35-Cdc42* GTPase signal in order to regulate Wnt signalling upstream of *Arm*.

To determine whether JNK signalling has a conserved role in vertebrates, we utilised mouse colon organoids with a truncating mutation in *Apc* (*Apc^trunc^*) generated with CRISPR technology. We seeded WT and *Apc^trunc^* organoids, treated them with either DMSO or the JNK1 inhibitor DB07268 and extracted protein or measured proliferation rate (Fig. 6h). First, we observed that *Apc^trunc^* organoids displayed elevated levels of β-catenin resulting from its stabilisation upon activation of the Wnt pathway (Fig. 6i, j) [62]. Second, using a phospho-SAPK/JNK (Thr183/Tyr185) antibody, we find that *Apc^trunc^* organoids are in a JNK active state which could be suppressed by using DB07268 (Fig. 6i, k). Third, treatment with DB07268 preferentially reduced the proliferation of rate of *Apc^trunc^*organoids and had a mild non-significant effect on WT organoids (Fig. 6l, m). These findings demonstrate that JNK signalling is a conserved modulator of mouse colon organoid proliferation when *Apc* is mutated.

## DISCUSSION

In this study, we identified Rab35 as a negative regulator of progenitor proliferation and Wnt signalling when *Apc* is depleted. We show that Rab35 exerts this function by localising and activating Cdc42 on the plasma membrane. Consequently, Cdc42 potentiates JNK signalling which modulates the Wnt pathway upstream of β-catenin. We show that normal control over these processes is an important factor in maintaining progenitor proliferation, differentiation and for the longevity of flies. In *Apc* mutant mouse colon organoids, we demonstrate that maintenance of JNK signalling is essential for the proliferation of cancer organoids.

### Rab35 activity and localisation impacts intestinal function

Rab GTPases are reported to carry out their functions by interacting with a variety of effector proteins when they are active [63]. Our findings reveal that the loss of Rab35 function leads to Cdc42 activation and its translocation to the plasma membrane. Subsequently, Cdc42 acts as a molecular switch to relay the activity status of Rab35 in order to positively regulate JNK signalling. This is surprising given that there is limited knowledge on how GDP bound Rabs lead to the activation of proteins. [64]. One class of proteins that may be interesting to explore in the future are the Cdc42 GEFs. Previous reports indicate that mutations to *Apc* stimulate Asef, a Cdc42 GEF, by relieving its autoinhibition [65]. Asef contains a PH domain which mediates its PtdIns(3,4,5)P_3_-dependent membrane translocation. Interestingly, loss of Rab35 activity has been associated with an increase in PtdIns(3,4,5)P_2_ and previous reports indicate that Rab35 and PtdIns(3,4,5)P_3_ co-localise at phagocytic cups [37, 66]. Thus, it may be that Rab35, through its modulation of PtdIns(3,4,5)P_2_ / PtdIns(3,4,5)P_3_, is able to recruit Asef which in turn activates Cdc42. Asef has been proposed to be involved in the fragmentation of Golgi [67]. Our electron micrographs indicate that loss of *Apc* or *Rab35* did not alter the Golgi architecture. However, it remains to be determined what role Rab35 activity has in regulating Golgi fragmentation and the importance of this process for intestinal homeostasis.

Rab8 has been shown to regulate Wnt signalling at the level of Lrp6 [68, 69]. Our data indicates that Rab35 can control the level of membrane Arm downstream of *Arr/Lrp6* and *Apc* and has tumour suppressive roles in the intestine, contrary to its reported oncogenic properties *in vitro*[70]. Since Rab35 and Arm co-localise on ISC plasma membrane, are within a 40nm range, and Rab35 knockdown results in heterologous expression of hTfR, it is tempting to speculate whether Rab35 directly alters the trafficking of Arm or if membrane Arm reflects Wnt induced stabilisation of Arm. Interestingly, Rab11 and the exocyst components Sec5, Sec6 and Sec15 are involved in regulating the proper localisation of E-Cadherin and Arm [71]. Recently, other GTPases such as RalA/B have been reported to regulate Wnt signalling upstream of β-catenin [72]. Thus, further exploring the roles of GTPases in the intestine may uncover novel pathways that enable us to better understand Wnt regulation and diseases associated with this cascade.

### Regional Wnt activation in the intestine

The intestine is regionalised at the level of cell type, gene expression, morphology, pH and microbial populations [25, 73–75]. These differences persist in region-derived intestinal organoid cultures suggesting that location-specific functions are intrinsically programmed [76]. However, the regulation of regionalisation in the intestine has remained poorly understood, especially in the context of disease. Our study shows that the anterior region of the midgut has a particular susceptibility to Wnt perturbations. For example, silencing *Apc* in either all progenitors or ECs results predominantly in anterior Wnt activation. Importantly we found that Rab35 modulates regional Wnt activity, highlighting a key regulator during this process. Therefore, it is likely that there are intrinsic differences between the anterior and posterior intestine. It is worth noting that *Apc1* null mutant clones cause Wnt activation throughout the intestine [30], which may be a result of manipulating multiple cell types simultaneously as opposed to just progenitor cells. Interestingly, regional differences between proximal and distal CRC results in distinct clinical outcome for patients which is a poorly understood phenomenon [77, 78]. By using *Drosophila* as a genetic model to better understand the regional characteristics of CRC, we may in the future identify pathways that lead to the development of targeted therapies.

One surprising observation is that expressing *Apc^RNAi^*or *Arm^CA^* does not produce the same Wnt activation pattern in the midgut. While *Apc^RNAi^* increased Fz3-RFP in some ISCs and ECs, expression of *Arm^CA^* induced a predominantly cell autonomous response. These differences may point towards additional functions for *Apc* in activating Wnt signalling which are not shared by Arm or it may alternatively be due to the different strengths of transgenic construct used. Our *in vivo* clonal analysis also demonstrates that *Apc^RNAi^* clones respond non-uniformly to Wnt transduction. Therefore, this suggests that Wnt signalling may be stochastic and/or dependent on cellular state, as has been previously suggested in the leech embryo [79].

### Blackbelt as a putative GAP

Our single cell analysis of the intestine revealed that *blackbelt* (*CG42795*) is overexpressed in progenitor cells when *Apc* is silenced by *RNAi*. Blackbelt is predicted to enable GTPase activator activity. The human orthologue of Blackbelt is TBC1D30 which has been reported to display Rab8 GAP activity *in vitro* [80]. It is generally assumed that GAP proteins normally have one Rab substrate. However, previous studies have shown that this is not the case, and a single GAP can stimulate GTPase activity of several Rabs [81]. Interestingly, in addition to the Rab-GAP-TBC domain, Blackbelt contains a large extension at the C-terminus that is conserved only in *Drosophila*. Thus, it is feasible that Blackbelt may have additional substrates in flies. Our structural modelling suggests that Blackbelt is a conventional GAP containing R and Q finger that can interact with Rab35. However, it remains to be seen if this interaction is present *in vivo* and if Blackbelt can stimulate Rab35 GTPase activity.

### Cross talk between Rab and Rho GTPases in the intestine

Our data shows that Rab35 modulates Wnt activity by regulating Cdc42 and JNK signalling, which are components used by non-canonical Wnt signalling to exert downstream functions. Rho GTPases have been implicated in canonical Wnt signalling in the past. For example, activation of Rac1 controls nuclear localisation of β-catenin during canonical Wnt signalling and is required for intestinal tumorigenesis [82, 83]. Moreover, JNK has been reported to directly phosphorylate β-catenin and JNK activation is associated with intestinal progenitor proliferation [82, 84–86]. Cdc42 has also been implicated in the turnover of β-catenin in skin and is able to directly bind Apc [87, 88]. Our findings provide *in vivo* mechanistic insight into the link between Cdc42 and JNK signalling and its role in canonical Wnt regulation when Apc is non-functional.

Developing effective Wnt pathway inhibitors has proved challenging given the toxicity associated with targeting a key homeostatic pathway. Interestingly, we show that neither Cdc42 nor JNK signalling impacts the Wnt pathway during homeostasis. Indeed, *Cdc42* is highly expressed in colorectal adenocarcinoma which provides an incentive to develop inhibitors for this protein as a form of therapeutic treatment [89, 90]. Moreover, our experiments in mouse colon organoids suggest that maintaining active JNK signalling is important for the proliferative capacity of stem cells and that blocking this preferentially restrains *Apc^KO^* organoid growth. Thus, our study highlights the importance of maintaining active JNK signalling in cancer organoids and provides a mechanism that can curb Wnt activated cells with minimal impact on wild type cells.

## MATERIALS AND METHODS

### Fly husbandry

*Drosophila* stocks were raised on a 12:12 hour light:dark cycle and maintained on standard fly food consisting per litre of 44 g syrup, 80 g malt, 80 g corn flour, 10 g soy flour 18 g yeast, 2.4 g methly-4-hyroxybenzoate, 6.6 mL propionic acid, 0.66 mL phosphoric acid and 8 g agar. For experiments requiring temperature shift (*Gal80^TS^*) for transgene induction, parental lines were kept at 18°C and the progeny were shifted to 29°C after eclosion for 20 days. For experiments where temperature shift was not required, flies were kept at 29°C to ensure age-dependent factors do not confound comparison between experiments. For CRISPR mutagenesis experiments, newly eclosed flies were shifted to 29°C for 10 days and subsequently to 18°C for 30 days before switching back to 29°C for one day prior to imaging as previously reported [91]. For all experiments, mated female flies were used and transferred to fresh food every two days.

### Fly stocks

The following fly lines were used in this study: *esg^TS^, UAS-GFP* (gift from B. Edgar [27])*, esg^TS^*(gift from B. Edgar), *esg^TS^, Su(H)GBE-Gal80, UAS-GFP* (gift from B. Edgar)*, Su(H)GBE^TS^, UAS-GFP* (gift from B. Edgar [92])*, Myo1a^TS^, UAS-GFP* (gift from J. Zhou [36])*, Rab35-Gal4 (BL 51599), esg^TS^, UAS-H2B-RFP* (gift from Tobias Reiff [44])*, esg^F/^*^O^ (gift from P. Patel [36])*, esg^TS^, UAS-GFP, puc-LacZ* (gift from J. Zhou)*, esg^TS^, UAS-GFP, UAS-Cas9^p.2^* (gift from F. Port)*, Act-Cas9, Tub-Gl4* [93]*, hh-Gal4* [94]*, Fz3-RFP* (gift from Y. Ahmed)*, UAS-Golgi-RFP (BL 30907), Rab35^CRISPR.GFP^*(gift from J.T. Blankenship)*, UAS-hTfR-GFP* (BL 36858)*, sqh-ChFP-Cdc42* (BL 42237)*, Arm-GFP* (BL 8555)*, UAS-Apc^RNAi^* (VDRC 51469, gift from S. Hirabayashi*), UAS-Rab35^RNAi^* (BL 80457)*, UAS-Rab35^DN^*(BL 9819), *UAS-Rab35^CA^* (BL 9818)*, UAS-Rab35^WT^*(BL 9821)*, UAS-wg^RNAi^* (VDRC 39670)*, UAS-Evi^RNAi^*(VDRC 5214)*, UAS-Pan^DN^* (BL 4785)*, UAS-blackbelt^RNAi^*(VDRC 108779)*, sqh-WASp.RBD-GFP* (BL 56746)*, UAS-Cdc42^WT^* (BL 28873)*, UAS-Cdc42^DN^* (BL 6286)*, UAS-Cdc42^RNAi^* (BL 28021)*, UAS-Bsk^DN^* (BL 9311)*, UAS-puc^RNAi^* (BL 34392)*, UAS-Hep^RNAi^* (BL 35210), *UAS-arr^HA^* (BL 42691)*, UAS-Ras^V12^* (gift from S. Hirabayashi*), UAS-N^RNAi^* (VDRC 27228)*, UAS-Rab4^RNAi^* (BL 33757)*, UAS-Rab5^RNAi^* (VDRC 34096)*, UAS-Rab11^RNAi^* (BL 27730)*, UAS-Arm^CA^* (BL 4782)*, UAS-Apc1^gRNAx2^* (This study)*, UAS-Apc1+2^gRNA^* (This study), *UAS-Rab35^gRNAx2^*(This study)*, UAS-blackbelt^gRNAx2^* (This study).

### sgRNA generation

Transgenic flies expressing Cas9 sgRNAs were generated as previously described [95]. Briefly, PCR amplicons containing spacer sequences for two sgRNAs were generated from plasmid pCFD6 (Addgene 73915) using primers sgRNAampfwd: CatGAAGACCTTGCANNNNGTTTCAGAGCTATGCTGGAAAC and sgRNA amprev: CatGAAGACCCAAACNNNNTGCACCAGCCGGGAATCGAACC (were NNNN denotes the spacer sequence or reverse complement thereof). Spacer sequences were TGAGTAAACACACTCCACCT and CATGAGGCCGTAGCTCCTGA for the lines targeting APC1 and APC2, CCACCTCGGATTACAACCAG and GGGCGAGGTTAGGCGGGCAC for Blackbelt, CTGCAGATCTGGGACACGGC and GATCCAGAATAACTGCGATG for Rab35 and GTCGTCCAGGCAGCTCCCTT and CAGGTTCCATAGAGTTCCAC for APC1. Amplicons were generated with the Q5 2x mastermix (New England Biolabs) and cloned by GoldenGate assembly into pCFD5w (Addgene 112645; APC1 and Rab35) or pCFD6 (APC1/2 and Blackbelt). Expression plasmids were integrated into the attP40 landing site in the *Drosophila* genome using standard injection procedures.

### Genetic modifier screen

Candidate genes (*UAS-x*) with membrane trafficking functions were selected based on their misexpression in colorectal cancer patients or their prior association with regulating signalling pathways. A modifier screen was performed by crossing *UAS-x* lines for these candidate genes to a parental line which stably expressed *Apc^RNAi^* using the *esg^TS^* system. 1-3 day old progeny (*esg^TS^> Apc^RNAi^* + *UAS-x*) were shifted to 29°C for 20 days and the intestine was dissected and imaged. A scoring system was developed based on the number of progenitor cells throughout the intestine, where paired interactions between *Apc^RNAi^* + *UAS-x* were compared to *Apc^RNAi^* only. Scores ranged from -2 to +2, where negative scores indicate that paired interactions reduced progenitor cell numbers and positive scores indicate that paired interactions enhanced progenitor cell number when compared to *APC^RNAi^*. Fly lines that were included in the modifier screen are listed in Supplementary table 1.

### Escargot-Flip-Out experiments

Flip-out clones were generated as previously described [36]. Briefly, expression of *flippase* by esg^TS^ at 29°C activates a constitutive Act>STOP>Gal4 driver by excising the STOP cassette flanked by FRT sites. This system was induced for 10-15 days and results in expression of GFP and *RNAi* in both progenitor cells and their descendant progeny. Flies were evenly housed in control and treatment groups and their food was changed every 2 days throughout the experiment.

### Dissection and immunohistochemistry

Flies were starved for 3h prior to dissection to reduce luminal content in the intestine. Adult female intestines were dissected in PBS (Phosphate buffered saline, P3812-10PAK) and transferred to Polylysine slides and fixed in 4% Paraformaldehyde (16 % Paraformaldehyde (Thermo Scientific) diluted in PBS) for 20-60 min depending on the antibody. Samples were washed with PBST (PBS with 0.1% Triton X-100) for 30 min and then blocked with PBSTB (PBST with 1% Bovine serum albumin) for 30 min at room temperature. Primary antibody was diluted in PBSTB and incubated with samples overnight at 4°C. Samples were then washed five times in PBST and incubated for 1.5 h-2.5h at room temperature with secondary antibody (antibodies coupled to Alexa fluorophores, Invitrogen) in PBSTB. Samples were washed five times in PBST and mounted in mounting medium (VECTASHIELD from Vector Laboratories with or without DAPI; Vector Labs, H-1200 or H-1000 respectively). Immunostainings for both experimental and control conditions were carried out on the same slide to enable direct comparisons. The region of interest imaged is indicated in the figure legends. The following antibodies were used: mouse anti-Armadillo (1:50; DSHB, N27A1), rabbit anti-GFP (1:1000; Invitrogen, A-11122), mouse anti-Prospero (1:20; DSHB, MR1A), rabbit anti-β-Galactosidase (1:1000; ICL Lab, RGAL-45A), rabbit anti-Phospho-Histone H3 (1:500; Cell Signaling, 9701S). Conjugated fluorescent secondary antibodies (conjugated to AlexaFluor488, AlexaFluor549 and AlexaFluor549) were obtained from Invitrogen (Life Technologies) and used at 1:1000.

### Image acquisition and processing

Confocal fluorescent images were acquired using either an upright Nikon A1 confocal microscope with a 25x Apo dipping objective (NA 1.1) or a spinning disk microscope (CREST V3) on a Nikon Ti2 inverted microscope equipped with a 60x NA PlanApo 1.4 oil immersion objective using Nis-Elements 5.3 software. Images shown represent maximal intensity projection of stacks covering the first epithelial layer. The same acquisition settings (laser power and gain/camera settings) were applied to both experimental and control groups. All statistical analyses were performed on raw 16bit images using Fiji 2.0 (see detailed description below). For display, images were converted to 8bit using the same scaling within an experimental group. In some cases, the brighter signals are displayed beyond saturation in order to appreciate lower signals. To visualise the whole gut, images were stitched together using the Pairwise Stitching plugin in Fiji.

To enhance resolution in Figure 3b, we used a Nikon AX laser scanning confocal microscope with the N-SPARC avalanche detector in super resolution mode. Stacks of 4-5 µm were acquired with a 60x silicone immersion objective with NA 1.3 and deconvolved using the Richardson-Lucy Algorithm as part of the Nis-Elements software.

To quantify RFP and GFP positive cell numbers, a Fiji macro was developed by Dr. Damir Krunic from the DKFZ imaging facility. This macro was used on z-projected images of either the upper or entire epithelial layers of the intestine. For each channel, the background was subtracted and the Gaussian Blur was applied to smooth the cell/area being measured. Threshold for each signal intensity was determined and a “Find Maxima” code was applied to define the centre of the area or cell. GFP and RFP signal that overlapped with the cell centre defined by the DAPI channel was quantified. The percentage of GFP or RFP cells was calculated as a measure of total DAPI cells. To quantify midgut mitosis, phospho-Histone H3 puncta that overlapped with DAPI nuclei were counted throughout the entire intestine using a Nikon A1 confocal. To quantify the colocalisation between Rab35 and the Golgi, z-stacks were taken of the midgut and GFP/RFP double positive puncta were manually counted and scored against only GFP.

### Armadillo membrane intensity

The plot profile tool in Fiji was used to quantify Armadillo (Arm) on the ISC-ISC membrane by measuring the intensity profile of a line drawn perpendicular to the membrane. The midpoint intensity was used for all ISCs after subtracting the background. Each data point represents one ISC and a total of 3 ISCs were measured per midgut, see Supplementary table 3 for n numbers. The same method was used to quantify the membrane localisation of Rab35.

### Proximity ligation assay

Flies were dissected and incubated with primary antibodies against the two proteins of interest as described in the immunohistochemistry section. Intestinal samples for negative control and for experiments were arranged on two distant parts of the same slide which were divided with silicon. Primary antibodies used were from the two different host species, mouse anti-Arm and rabbit anti-GFP. The next day, probes that conjugate with primary antibodies, PLA Probe Mouse MINUS (DUO82004, Sigma-Aldrich) and PLA Probe Rabbit PLUS (DUO82002, Sigma-Aldrich) were diluted 1:5 in antibody diluent (DUO82008, Sigma-Aldrich). Samples were washed three times with PBST and incubated with 60μl PLA probe solution in a pre-heated humidity chamber for 1h at 37°C. For probe ligation, the samples were washed in PBST and then incubated with ligation solution containing in 5x Duolink Ligation buffer (DUO82009, Sigma-Aldrich) and Duolink Ligase (DUO82029, Sigma-Aldrich) in a pre-heated humidity chamber for 30 min at 37 °C. For amplification, the samples were washed in PBST and incubated with amplification solution containing in 5x Duolink Amplification Red buffer (DUO82011, Sigma-Aldrich) and Duolink Polymerase (DUO82030, Sigma-Aldrich) in a pre-heated humidity chamber for 30 min at 37 °C. Samples were then washed and mounted in mounting medium as described in Immunohistochemistry. The signal for the proximity of the proteins was observed in the red channel with Nikon A1 confocal.

### Cdc42 biosensor measurements

To measure GTP-bound Cdc42, we used a biosensor consisting of a RhoA binding domain of WASp fused to GFP under control of the sqh promoter. A cell profile was drawn around each ISC and the activity of GFP, which corresponds to CDC42^GTP^, was measured as mean grey value relative to ISC size after subtracting the background. Signal was present in the cytoplasm and nucleus. A total of 3 ISCs per midgut were measured, see Supplementary table 3 for n numbers for n numbers.

### Regional Wnt activity profiles

Fluorescence intensity was measured along three lines of defined position (‘upper quarter’, ‘middle’, and ‘lower quarter’) spanning the length of the entire midgut using the ImageJ plot profile tool from the MHB to the proventriculus as previously described. The average between the three lines was calculated and the data was fitted using the R (version 4.2.2) Local Polynomial Regression Fitting (loess) smoothing function with formula y ∼ x and the grey area around the lines representing the confidence intervals. The geom_smooth function from ggplot2 version 3.4.2 was used for the fitting. The distance measurements have been scaled by dividing each measurement by the maximum distance. The same tool was used to measure Cdc42 and PLA intensity across a line drawn perpendicular to the progenitor cell membrane.

### Infection experiments

For oral infection, flies were starved for 4h and then kept at 29 °C overnight in vials with pre-cut Whatman Filter Paper disk containing 5% sucrose solution with and without concentrated bacteria. The sucrose solution with concentrated bacteria was made by centrifuging high-dose bacteria (OD600=200) at 3500 g for 15 min and suspending the pellets with 5% sucrose solution.

### Lifespan assay

10-15 adult female flies and five adult male flies were kept in each vial as described in Fly husbandry at 29°C. All lifespan experiments started with at least 50 adult female flies for each condition and the number of dead female flies was recorded every day until all female flies were dead.

### Intestinal barrier assay

Experimental flies were starved for 3h and then transferred on to food containing 1% FCF blue dye and allowed to feed overnight. The next day, flies were washed in PBS and imaged using a Leica DFC420C camera attached to a Leica M165FC stereo microscope.

### Electron microscopy

Midguts from *Drosophila* flies were dissected in PBS and immediately immersed in EM-fixative (2.5% Glutaraldehyde, 1mM MgCl_2_, 1mM MCaCl_2_, in 100mM Ca-Cacodylate, pH 7.2). After primary fixation, further steps included post fixation in buffered 1% osmium tetroxide, en-bloc fixation with uranyl-acetate, graded dehydration with ethanol and resin-embedding in epoxide (Glycidether, NMA, DDSA: Serva, Heidelberg, Germany). Midguts were embedded and arranged longitudinally for sectioning. Ultrathin sections at nominal thickness 50 nm and contrast-stained with lead-citrate and uranyl-acetate were observed in a Zeiss EM 910 at 80kV (Carl Zeiss, Oberkochen, Germany) and micrographs taken using a slow-scan CCD-camera (TRS, Moorenweis, Germany).

### Adult *Drosophila* wing disc analysis

Fly wings were imaged by placing whole flies with the correct genotype in 1:1 glycerol/ethanol mixture for several hours until fully perfused. Afterwards, the wings were separated from the fly abdomen at the joint and mounted onto a microscope slide in glycerol/ethanol. The wings were imaged with the Cell Observer microscope (Zeiss) under brightfield illumination with the 2.5x objective.

### *Blackbelt* protein domain prediction

The domains of CG42795 were predicted using Pfam for *Drosophila melanogaster*, and complementary regions for other species were determined by checking alignment. A conserved TBC-GAP domain was highlighted in all species but *Drosophila* also contained an extended conserved C-terminal section which was considerably shorter in other species.

### Structural prediction

Structure of CG42795 was predicted using AlphaFold2.0 via Cosmic2 [96]. Five models were generated and based on the pLDDT score, all are similarly reliable. It has to be noted, that the long Drosophila specific gene extension is mostly unstructured and no prediction was possible to make for this region. Only the Rab-Gap-domain and the TBC1-domain regions are reliable above 80% pLDDT. The protein-protein interaction between Rab35 and CG4759 was predicted using ColabFold on the Cosmic2 [97]. All five predicted models have a similar IDDT score and again only structural predictions for conserved domain regions are significant. Models were analysed and visualised using PyMOL Version 2.4.0.

### Phylogenetic analysis

Sequences for all used species were obtained from Flybase or NCBI databases. Protein sequences of CG42795 were aligned using ClustralO implemented in Seaview Version 5.0.5. The obtained alignment was trimmed by hand and saved as a phylip file for further analysis. Maximum likelihood tree was created using IQtree 2.1.2 with classical bootstrapping and UltraFast boostrapping (Total number of iterations: 102). Substitution model was predicted using the Model finder function in IQtree. The model JTT+F+R4 was predicted as the best fitting model. Tree visualisation was done in Seaview and organism pictures are from Phylopic.

### Single cell RNA-sequencing and high-throughput sequencing

For scRNA-seq, flies were starved for 3h and placed in vials with filter paper containing 5% sucrose for 16h. 20 midguts were dissected from respective genotype in ice-cold PBS, taking care to remove the hindgut, Malpighian tubules and proventriculus. Samples were digested in 1 mg/ml Elastase (Sigma, #E0258) solution at 25°C for 45 min at 1000RPM and vortexed every 15min. Samples were then pelleted and resuspended in PBS and dissociated cells were then passed through a 40uM than 20uM cell strainer and subsequently counted. Approximately 20,000 live cells were used for scRNA-seq with 10X Genomics 5’ V2 kit following the manufacturers protocol for library generation. Prior to sequencing, library fragment size was determined using an Agilent Bioanalyzer high-sensitivity chip and quantified using Qubit. Libraries were multiplexed and sequenced using a Nextseq 550 at the Deep Sequencing Facility, BioQuant, Heidelberg University.

### Single cell RNA-sequencing data analysis

Single-cell RNA-seq data quality control and analysis were conducted using the R package Seurat (version 4.3.0). Cells with reads of fewer than 200 genes were filtered out, and genes expressed in fewer than 3 cells were also excluded. Expression values were log-normalised before visualisation. For UMAP visualisations, replicates and conditions were integrated using the Seurat function IntegrateData, followed by Principal Component Analysis and Louvain clustering. Cell types were identified using automatic reference-based annotation from the Seurat package, with the flycell atlas data set as the reference [51, 52]. Cells that could not be automatically annotated were manually assigned using marker genes (Extended Data Fig. 6B) for expected cell types. For differential expression analysis, cell types were aggregated into pseudo-bulks by summing up counts for each cell type. Differential expression analysis between conditions was performed on the pseudo-bulk counts using the R package DeSeq2 (version 1.38.3 [98]) with the Wald test and accounting for different batches. Gene set enrichment analysis was carried out using the R package clusterProfiler (version 4.6.2 [99]) on genes ranked by their Wald statistic from the differential expression analysis.

### Organoid culture

Mouse colon organoids were generated as described by [100]. APC mutations were induced by transfection of “wild-type” organoids with the PX459, pSpCas9(BB)-2A-Puro, plasmid containing a sgRNA targeting APC (5‘-GGCACTCAAAACGCTTTTGA-3‘) using lipofectamin-2000 (Thermo Fisher Scientific, 11668027). WT organoids were grown in WENR medium and APC truncated organoids were grown in ENR medium and are later refererred to as mCol WT and mCol APC. Organoids were grown in BME Type R1 (Cultrex, 3433-010-R1) and split every 6-7 days. ENR consists of 70% v/v of Advanced/DMEM F12 (Life Technologies, 12634028) supplemented with 1% v/v penicillin/streptomycin solution (Life Technologies, 15140122), 1% v/v HEPES buffer (Sigma, H0887) and 1% v/v Glutamax (Life Technologies, 35050038), supplemented with 10% Noggin-conditioned medium, 20% R-spondin1-FC-conditioned medium, 1x B27 (Life Technologies, 17504044), n-Acetyl-cysteine (1.25 mM, Sigma, A9165), EGF (50 ng/ml, PeproTech, 315-09-1000), Y-27632 (10 μM, Selleck Chemicals Co., SEL-S1049) and Primocin (100 μg/ml, InvivoGen, ant-pm-2). WENR is ENR medium supplemented with 5 nM Wnt-surrogate (Life Technologies, PHG0402). For experiments, organoids were seeded at a density of 750-1000 cells/µl. 72 hours after seeding, the organoids were treated with 10 uM DB07268 (Medchemexpress, HY-15737-10) for 72 hours. Control organoids were treated with equal volume of DMSO (Sigma, D8418). Images were taken on a Keyence BZ-X810 microscope.

### Western blot assay

Organoids were harvested in cell recovery solution (Corning, 11543560) to remove the extracellular matrix. Subsequently, organoids were lysed in RIPA buffer (Life Technologies, 89900) supplemented with cOmplete (TM) protease inhibitor (Roche, 4693159001) and PhosSTOP tablet (Roche, 4906837001). 20 µg protein was loaded on a 4-12% Bis-Tris Bolt gel (Life Technologies GmbH, NW04120BOX). Proteins were blotted on a PVDF membrane (Merck Millipore, IPVH00010). Membranes were incubated in primary antibodies mouse anti-Vinculin (Sigma, V9264, 1:10 000), mouse anti-beta-catenin (BD 610154, 1:1000), mouse anti-hospho-SAPK/JNK (Thr183/Tyr185, Cell Signaling 9255, 1:500), rabbit anti-SAPK/JNK (Cell signaling, 9252, 1:500). Membranes were developed using SuperSignal West Femto Maximum Sensitivity Substrate (Life Technologies GmbH,34095), Immobilon Western HRP Substrate (Millipore, WBKLS0100) or fluorescent Starbright Blue700 secondary antibodies (Biorad,12004161) and bands were imaged on a BioRad ChemiDoc MP (BioRad,12003154). Quantification of bands was performed using the ImageLab 6.1 software (BioRad).

### Cell viability assay

Organoid growth was determined by using the CellTiter-Glo Luminescent Cell Viability Assay Kit (Promega, G7573) according to the manufacturers’ protocol, with minor modifications. On the day of readout, medium was removed and 100 µl of 1:1 mixture of CTG reagent and Advanced/DMEM F12 was added to the organoids. The plate was incubated for 30 min at room temperature, subsequently, 50 µl supernatant was transferred to a white lumino plate and readout on a Mithras LB940 plate reader (Berthold Technologies, Bad Wildbad, Germany).

### Statistics

GraphPad Prism 10 software was used for statistical analyses. Information on sample size and repetitions are listed in Supplementary table 3. The statistical tests used for each experiment are indicated in the figure legends.

## ACKNOWLEDGMENTS

We would like to thank B. Edgar, T. Blankenship, Y. Ahmed, P. R. Hiesinger, N. Tolwinski, N. Perrimon, J.P. Vincent, J. Cordero and P. Patel for providing fly lines. We would like to thank A.M.L Coenen-Stass and D. Ibberson for helping to identify which candidates to screen and for facilitating high-throughput sequencing at the Deep sequencing facility at Heidelberg University. We would like to thank the Nikon imaging facility and the Light and Electron Microscopy core facilities of DKFZ for support. We thank Cornelia Redel for providing helpful comments on an earlier version of this manuscript. We thank the Bloomington Drosophila Stock Center and the Vienna Drosophila Resource Center for supplying fly stocks.

## AUTHOR CONTRIBUTIONS

Conceptualization: S.R and M.B. Investigation: S.R., T.W., K.E.B., SR., S.B., F.P., B.P., R.D., N.R. Single cell RNA-sequencing: S.R., T.W., T.C., S.L., W.H. Electron microscopy: K.R. and S.R. Structural modelling and phylogenetic analysis: M.H. Super resolution imaging: T.W., S.R., U.E. Organoid experiments: K.E.B., S.R and S.R. Data analysis: S.R., T.W., E.V., U.E., T.C., W.H., M.B Supervision: M.B., W.H and S.R. Writing—original draft: S.R., T.W and M.B. Writing—review and editing: S.R., T.W., F.P and M.B, with input from all authors.

## DATA AND MATERIAL AVAILABILITY

All data needed to evaluate the conclusions in the paper are present in the paper and/or the Supplementary Materials. The described sgRNA strains are available from the corresponding authors. High-throughput sequencing data have been deposited at the EMBL-EBI European Nucleotide Archive project submission access PRJEB70885.

