## Extended Data Figure for "Restraining Wnt activation and intestinal tumorigenesis by a Rab35 dependent GTPase relay"

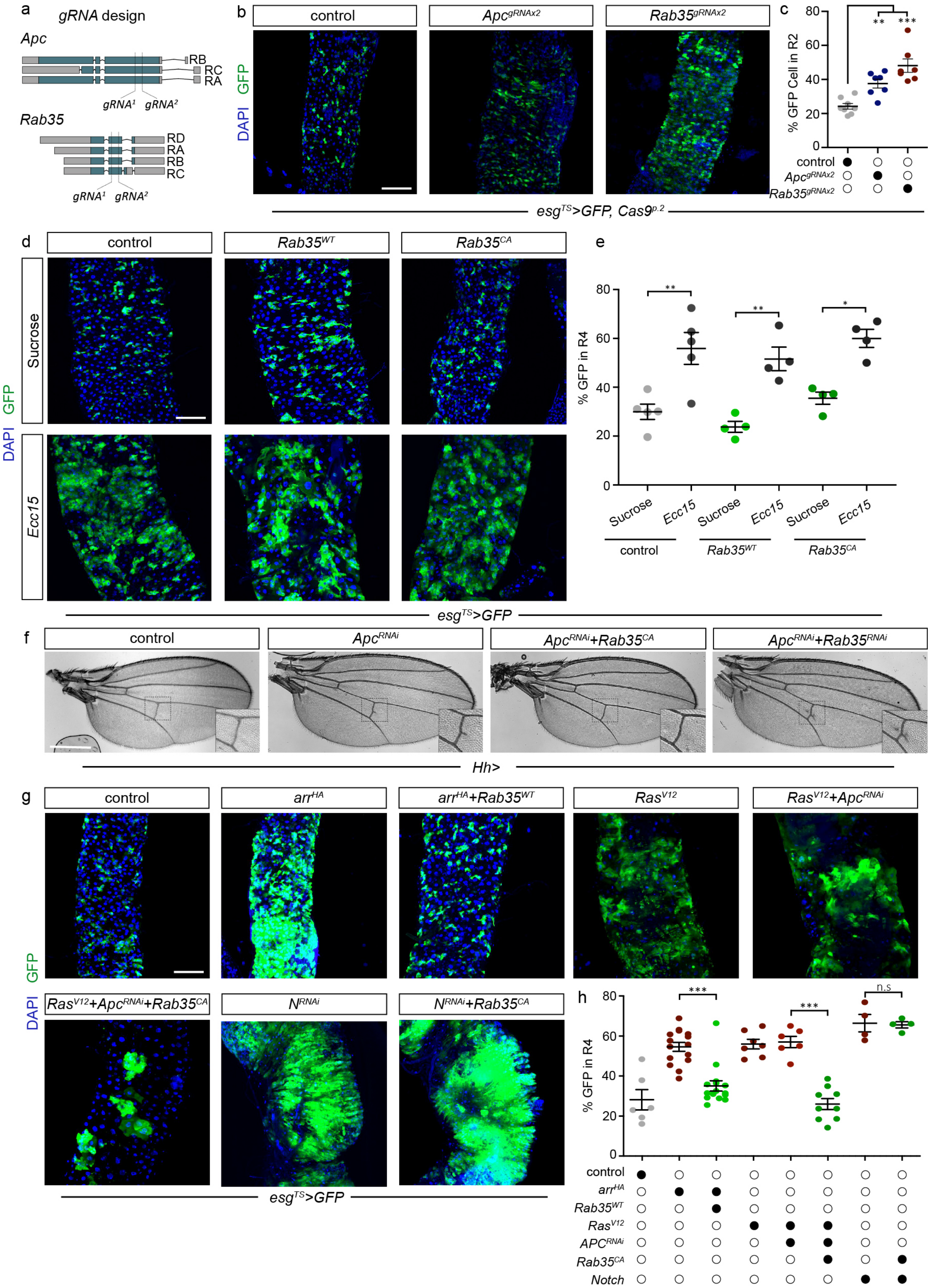


**Extended Data Figure 1**. a) Design of *gRNA* targeting two sites within the *Apc* and *Rab35* locus. b) CRISPR mediated mutagenesis of *Apc* and *Rab35* in progenitor cells increases progenitor proliferation. c) Quantification of *GFP*$^{+}$ cells in the R2 region. d) Intestinal response to pathogenic *Ecc15* infection is maintained when *Rab35* is either overexpressed (*Rab35*$^{WT}$) or constitutively activated (*Rab35*$^{CA}$) in progenitor cells using the *esg*$^{TS}$ driver. e) Quantification of *GFP*$^{+}$ cells in the R4 region. f) Knockdown of *Apc* in the wing disc using *hh-Gal4* results in ectopic vein formation at the posterior cross-vein section, while co-expression with *Rab35*$^{RNAi}$ enhances this phenotype and *Rab35*$^{CA}$ can rescue it. g) Genetic interaction of *Rab35* with various transgenes that induce proliferation of progenitor cells. *Rab35* can suppress progenitor cell proliferation in flies expressing *Arr*$^{HA}$ or *Ras*$^{V12}$+*Apc*$^{RNAi}$ but not *N*$^{RNAi}$ using the *esg*$^{TS}$ driver. h) Quantification of *GFP*$^{+}$ cells in the R4 region. Graphs represent the mean with standard error of the mean. One-way ANOVA test with Tukey post hoc comparison were used for all graphs. *$P < 0.05$, **$P < 0.01$. ***$P < 0.001$. Scale bar b, d, g 100 µm, f 500 µm.

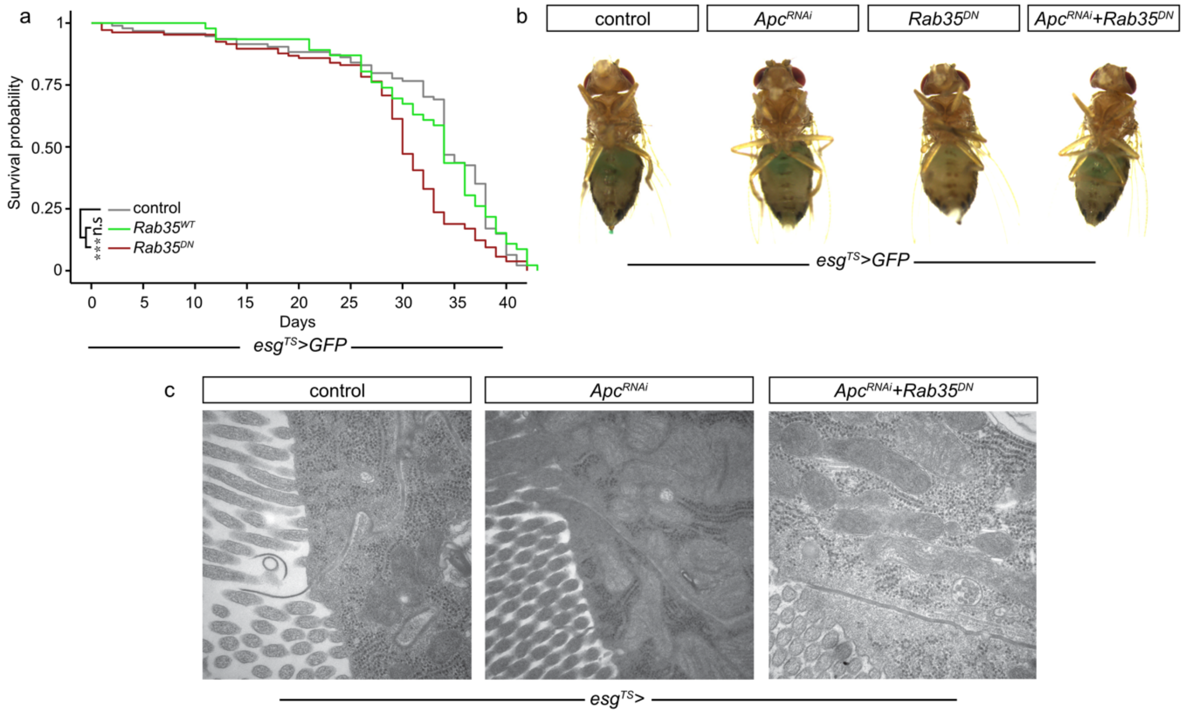


**Extended Data Figure 2**. a) Loss of *Rab35* activity (*Rab35*$^{DN}$) in progenitor cells using the *esg*$^{TS}$ driver decreases the lifespan of flies, while expressing a wild type form of *Rab35* (*Rab35*$^{WT}$) does not have a significant effect. b) Smurf assay was used to determine the integrity of the intestinal epithelium. None of the genetic perturbations influenced intestinal barrier integrity. c) Electron micrographs of the EC-EC junctions, demonstrating that they are intact in all conditions. A logrank test was used for lifespan experiment. *$P < 0.05$, **$P < 0.01$. ***$P < 0.001$.

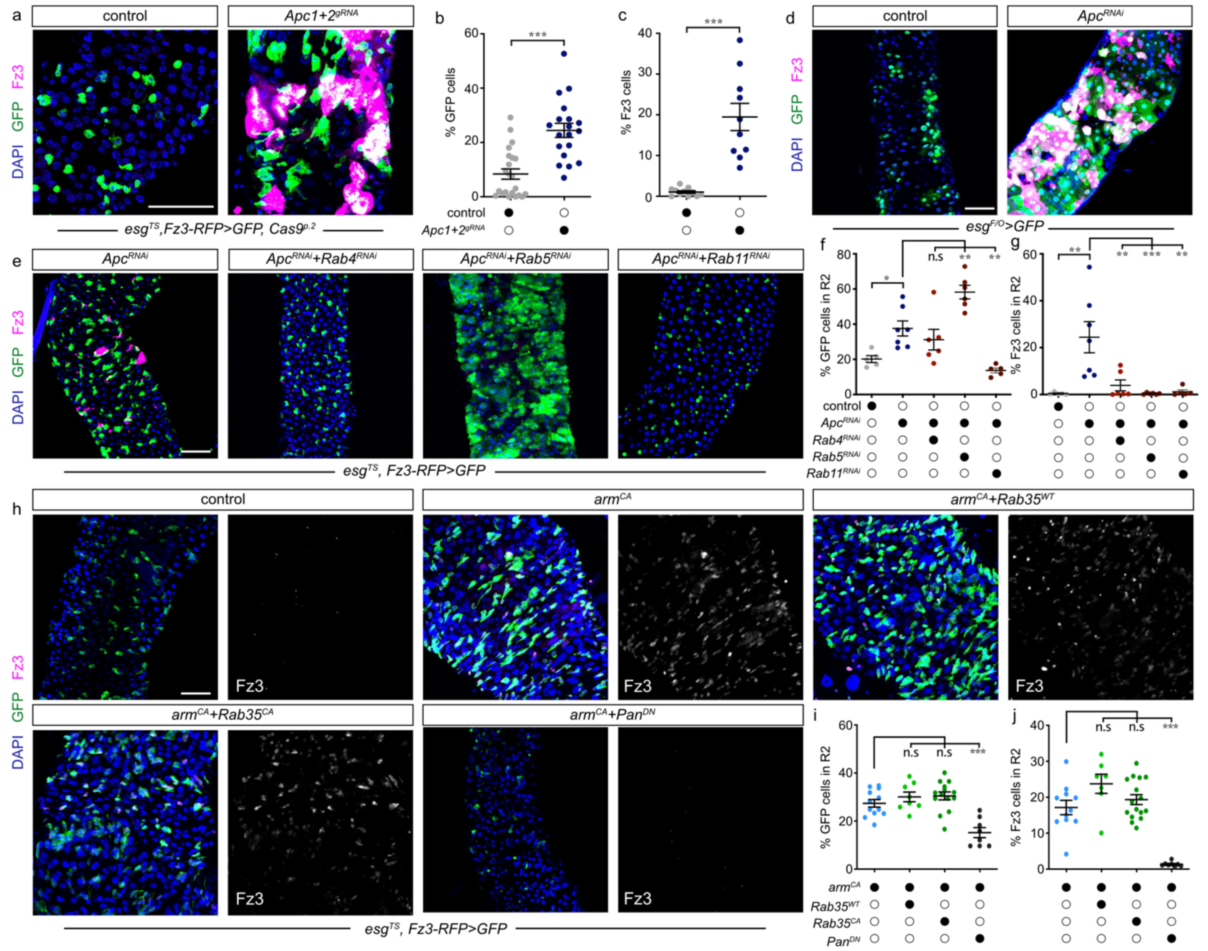

**Extended Data Figure 3**. a) CRISPR mutagenesis of *Apc1+2* results in widespread Wnt activation and progenitor proliferation in the R2 region using the *esg$^{TS}$* driver. b) Quantification of *GFP*$^{+}$ cells in R2 region. c) Quantification of Wnt activity (*Fz3-RFP*) in the R2 region. d) Generating flip out clones of *Apc$^{RNAi}$* results in stochastic Wnt activation in the intestinal epithelium. e) Knockdown of *Rab4* and *Rab11* decreases progenitor proliferation and Wnt activation when *Apc* is lost in progenitor cells, while *Rab5* knockdown enhances proliferation but not Wnt activity. f) Quantification of *GFP*$^{+}$ cells in R2. g) Quantification of Wnt activity (*Fz3-RFP*) in the R2 region. h) Expression of a constitutively active form of *Arm* (*Arm$^{CA}$*) using the *esg$^{TS}$* driver results in Wnt activation and progenitor proliferation, while expression of wild type or constitutively active *Rab35* does not supress this phenotype. Expression of a dominant negative form of *Pan* can rescue Wnt activity and progenitor proliferation when *Arm* is constitutively activated. i) Quantification of *GFP*$^{+}$ cells in R2 region. j) Quantification of Wnt activity (*Fz3-RFP*) in R2 region. Graphs represent the mean with standard error of the mean. One-way ANOVA test with Tukey post hoc comparison were used for all graphs except for b and c where a Mann-Whitney test was used. *$P < 0.05$, **$P < 0.01$, ***$P < 0.001$. Scale bar a, d, e, h 100 μm.

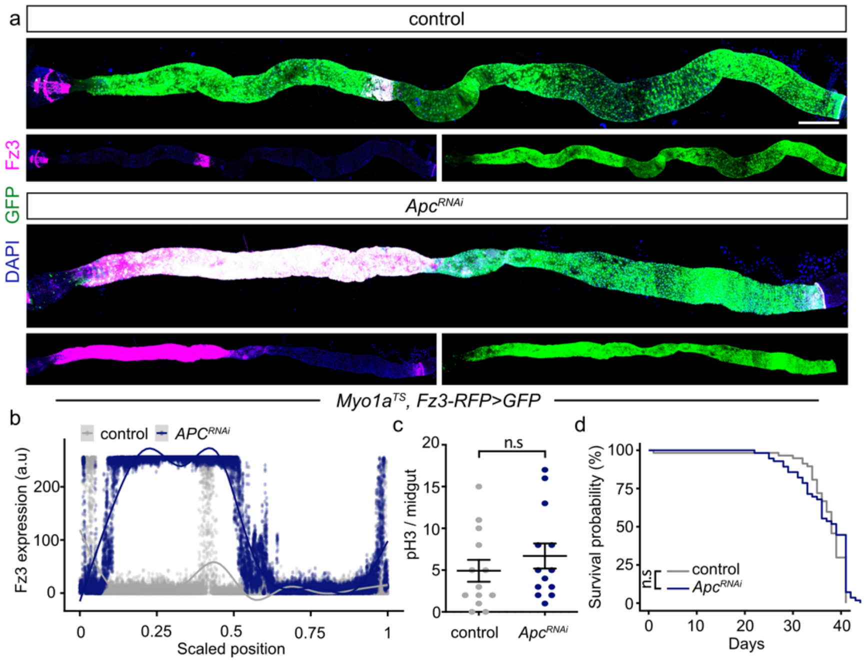


**Extended Data Figure 4**. a) Knockdown of *Apc* in all enterocytes using the *Myo1a*$^{TS}$ driver results in anterior Wnt activation. b) Quantification of *Fz3-RFP* along the intestinal epithelium. Scaled position 0 and 1 corresponds to the proventricular region and MHB boundary, respectively. c) Quantification of pH3 cells when *Apc* is knocked down in enterocytes using the *Myo1a*$^{TS}$ driver. d) Knockdown of *Apc* in enterocytes does not significantly affect the lifespan of flies. Scale bar a 200 µm.

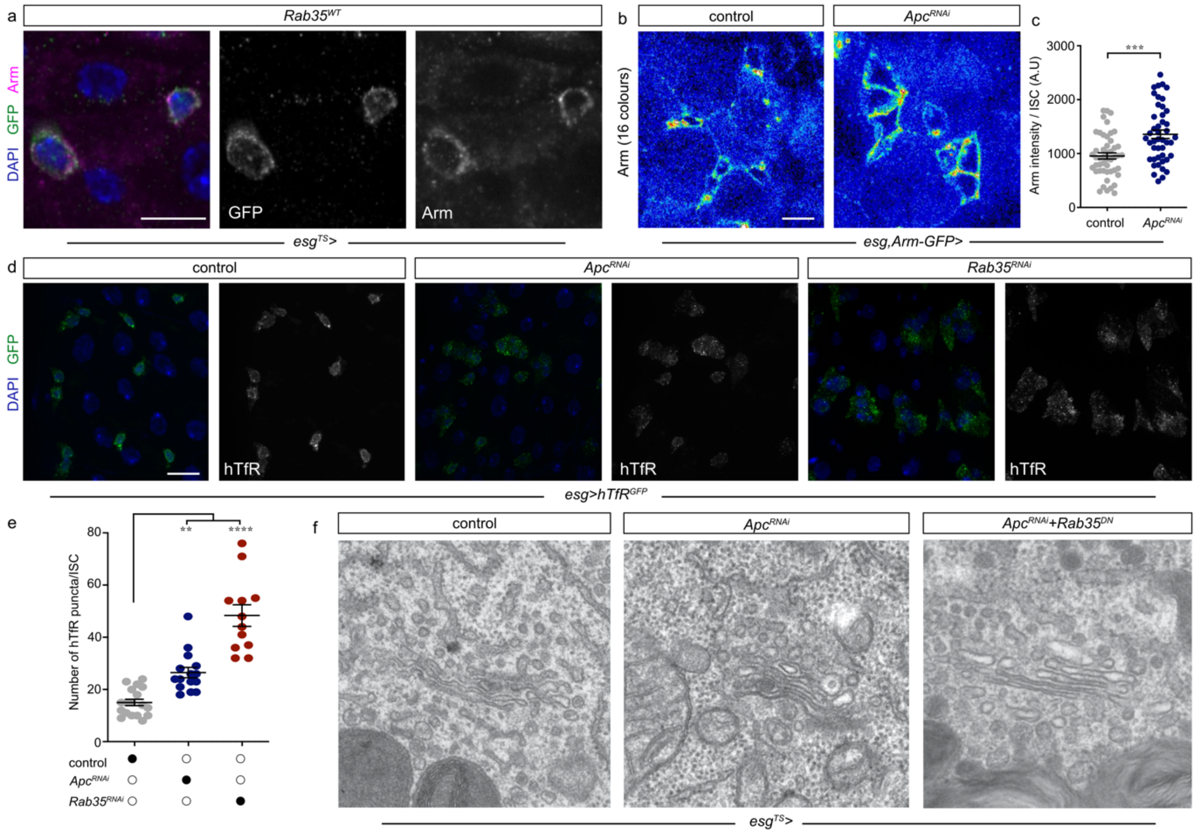


**Extended Data Figure 5**. a) Localisation of Arm and an overexpressed *GFP*-tagged wild type form of Rab35 (*$Rab35^{WT}$*) in progenitor cells. b) Knockdown of *Apc* increases endogenous Arm in progenitor cells. c) Quantification of endogenous membrane Arm when *Apc* is knocked down in progenitor cells. d) Expression of a *GFP*-tagged human transferrin receptor in progenitor cells highlights defects in its localisation when *Apc* or *Rab35* is knocked down. e) Quantification of human transferrin receptor in progenitor cells. f) Electron micrographs of the Golgi in progenitor cells illustrating no obvious morphological defects to the Golgi apparatus in different genetic backgrounds. Graphs represent the mean with standard error of the mean. A Mann-Whitney test was performed for panel C. One-way ANOVA test with Tukey post hoc comparison were used for panel e. *$P < 0.05$, **$P < 0.01$, ***$P < 0.001$. Scale bar a, b, d 10 µm.

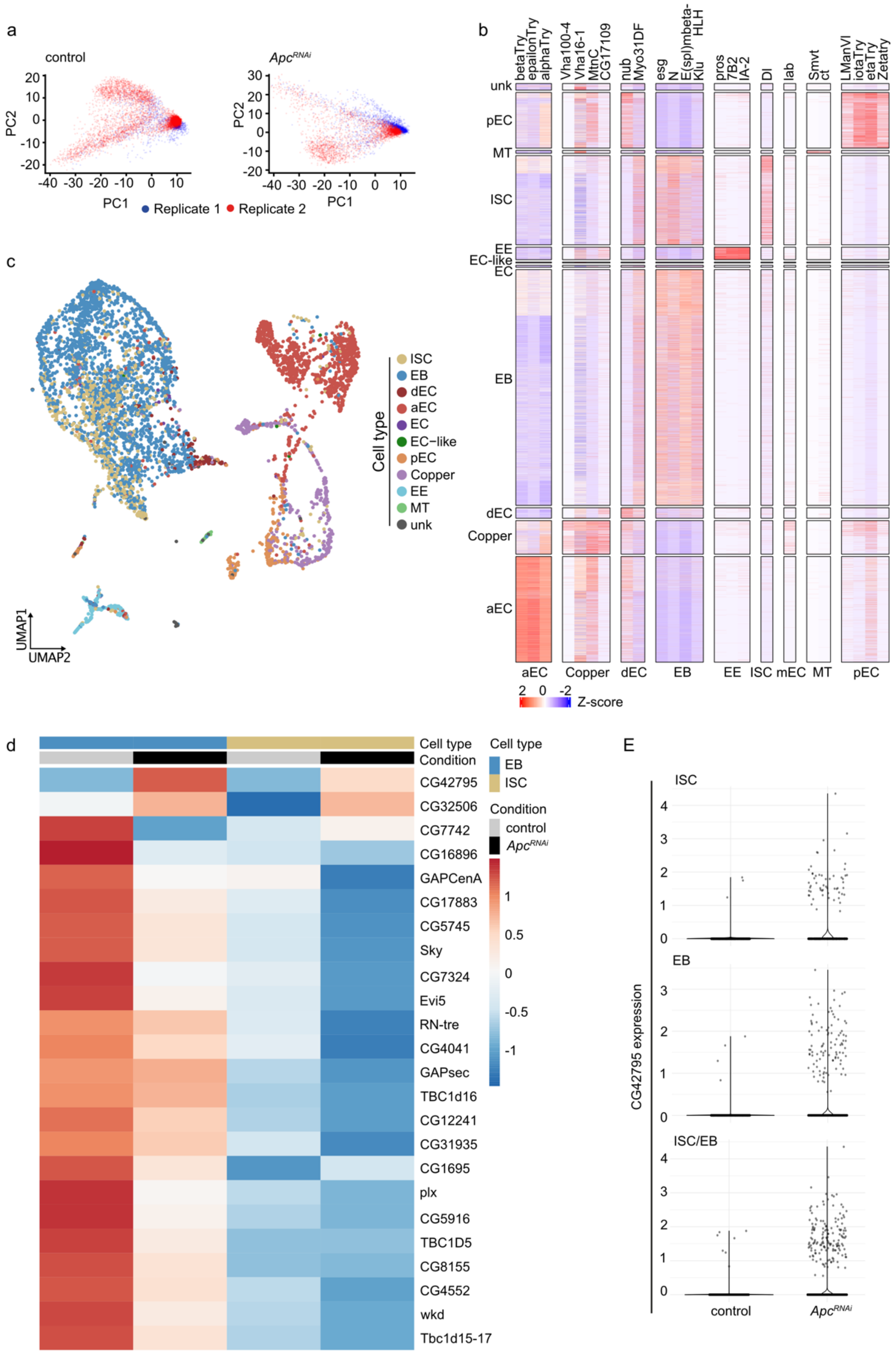


**Extended Data Figure 6**. a) PCA plot of scRNA-seq replicates under different conditions. b) Expression profiles of cell type specific markers used to annotate scRNA-seq datasets. c) UMAP of the intestine in *Apc RNAi* condition. d) Expression of Rab GAPs in different conditions and progenitor cell types across two replicates. *blackbelt* (*CG42795*) expression significantly increases in both ISCs and EBs when *Apc* is depleted. E) Volin plots displaying the expression of *blackbelt* in different progenitor cell types (ISC, EB, ISC/EB).

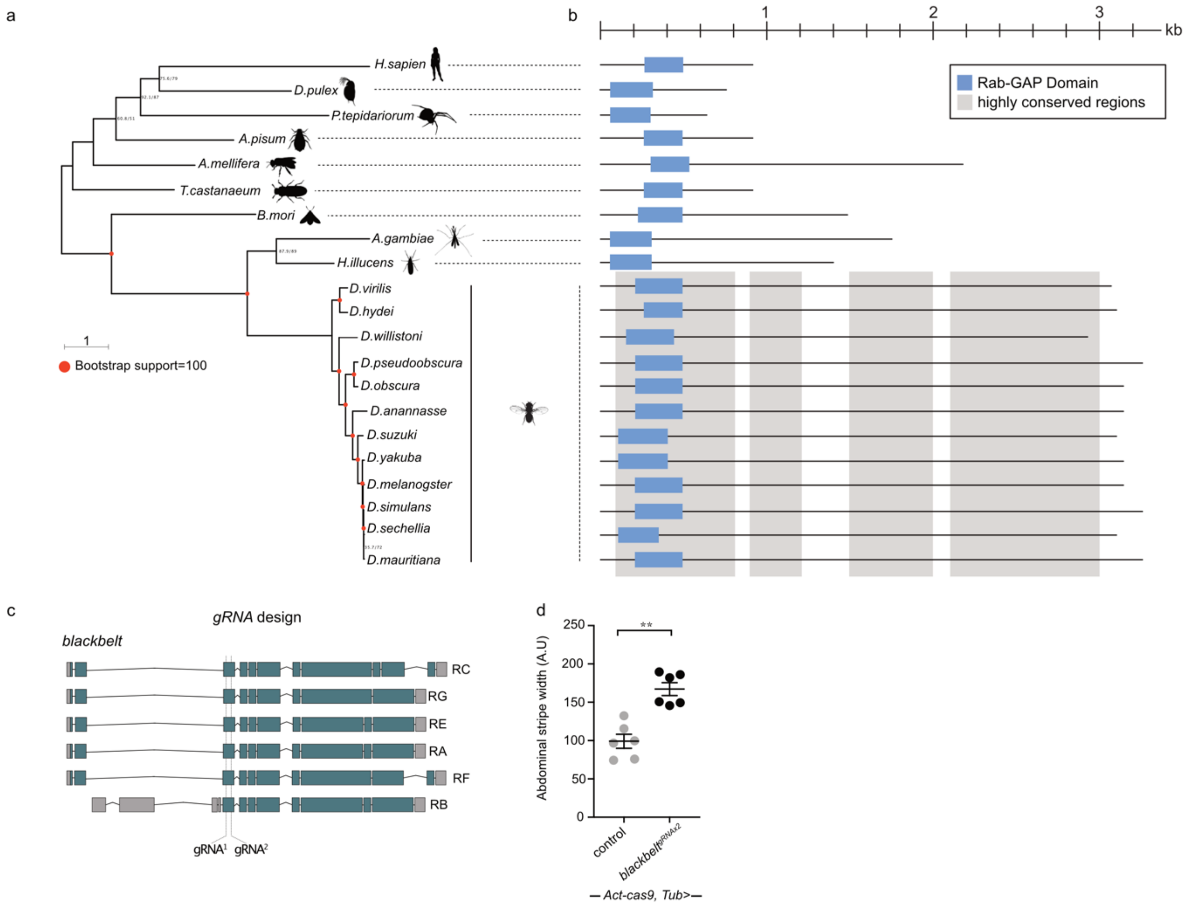


**Extended Data Figure 7**. a) Maximum likelihood tree showing the conservation of *blackbelt* within arthropods. *Homo sapiens* was used as outgroup. Bootstrap support is either shown at the nodes or depicted as red dot, if 100% support was predicted using IQtree. Left bootstrap value shows classic bootstrap predictions (SH-aLRT) and right value shows result from Ultrafast bootstrapping implemented in IQTree. Branch length indicates the nucleotide changes of the gene sequence. b) Schematic of Blackbelt domains in different species. Highly conserved regions in *Drosophila* are highlighted in light grey. Note, within the *Drosophila* species, Blackbelt protein size is larger than in other species while the GAP domain and the TBC1 domain are separated, unlike in other species that have their TBC1 domain is contained with the GAP domain. c) Design of gRNA targeting two sites within the *blackbelt* locus. d) Quantification of the width of A5 abdominal stripe in *blackbelt* mutants. Graphs represent the mean with standard error of the mean. A Mann-Whitney test was used for panel h. *$P < 0.05$, **$P < 0.01$, ***$P < 0.001$.

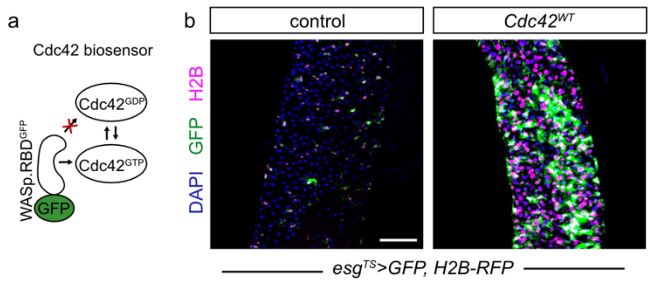


**Extended Data Figure 8**. a) Schematic of Cdc42 biosensor. Notice that WASp.RBD$^{GFP}$ only binds GTP and not GDP bound Cdc42. b) ReDDM lineage tracing system highlights that overexpression of wild type *Cdc42* increases progenitor proliferation and differentiation towards ECs. Scale bar a 100 µm.

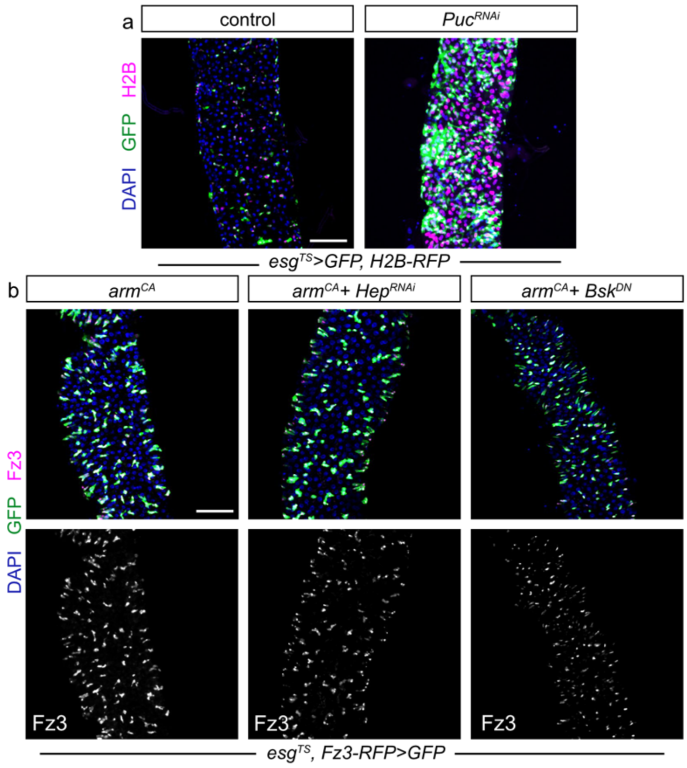


**Extended Data Figure 9**. a) ReDDM lineage tracing system reveals that activation of JNK signalling after expression of *Puc*$^{RNAi}$ results in accelerated proliferation and differentiation towards ECs. b) Blocking of JNK signalling by expressing *Hep*$^{RNAi}$ or *Bsk*$^{DN}$ cannot rescue Wnt activation or progenitor proliferation when *arm* is constitutively activated (*arm*$^{CA}$). Scale bar a and b 100 µm.
